# Post-translational modification by the Pgf glycosylation machinery modulates *Streptococcus mutans* physiology and virulence

**DOI:** 10.1101/2022.10.10.511621

**Authors:** Nicholas de Mojana di Cologna, Silke Andresen, Sandip Samaddar, Stephanie Archer-Hartmann, Tridib Ganguly, Jessica K. Kajfasz, Bruna A. Garcia, Irene Saengpet, Alexandra M. Peterson, Parastoo Azadi, Christine M. Szymanski, José A. Lemos, Jacqueline Abranches

**Affiliations:** Department of Oral Biology, University of Florida, College of Dentistry, Gainesville, FL, USA; Department of Microbiology, University of Georgia, Athens, GA, USA; Complex Carbohydrate Research Center, University of Georgia, Athens, GA, USA

**Keywords:** *Streptococcus mutans*, post-translational modifications, Cnm, glycosylation, phosphorylation, glycoproteomics, biofilm, competence, oral colonization

## Abstract

*Streptococcus mutans* is a keystone pathogen of dental caries, and the ability to form biofilms is essential for its pathogenicity. We identified a glycosylation machinery (Pgf) in *S. mutans* that post-translationally modifies two surface-associated adhesins, Cnm and WapA. The four *pgf* genes (*pgfS*, *pgfM1*, *pgfE,* and *pgfM2*) are part of *S. mutans* core genome and we hypothesized that the scope of Pgf goes beyond Cnm and WapA. By inactivating each *pgf* gene individually or creating a quadruple *pgf* mutant in *S. mutans* OMZ175, we showed that the Pgf machinery is important for biofilm formation. Compared to OMZ175, differences in surface charge, membrane stability, and genetic competence were also observed for most mutants. Importantly, *in silico* analyses and tunicamycin MIC assays suggest a functional redundancy between the Pgf machinery and the rhamnose-glucose polysaccharide synthesis pathway. Using a rat oral colonization model, we showed a 10-fold reduction in recovered CFUs for the *pgf* quadruple mutant compared to OMZ175. Finally, using Cnm as a model, we showed by glycoproteomics analyses that Cnm is heavily modified with N-acetyl hexosamine in OMZ175 whereas phosphorylations were observed for the *pgfS* mutant. Our findings indicate that the Pgf machinery participates in important aspects of *S. mutans* pathobiology.

**Graphical Abstract:** 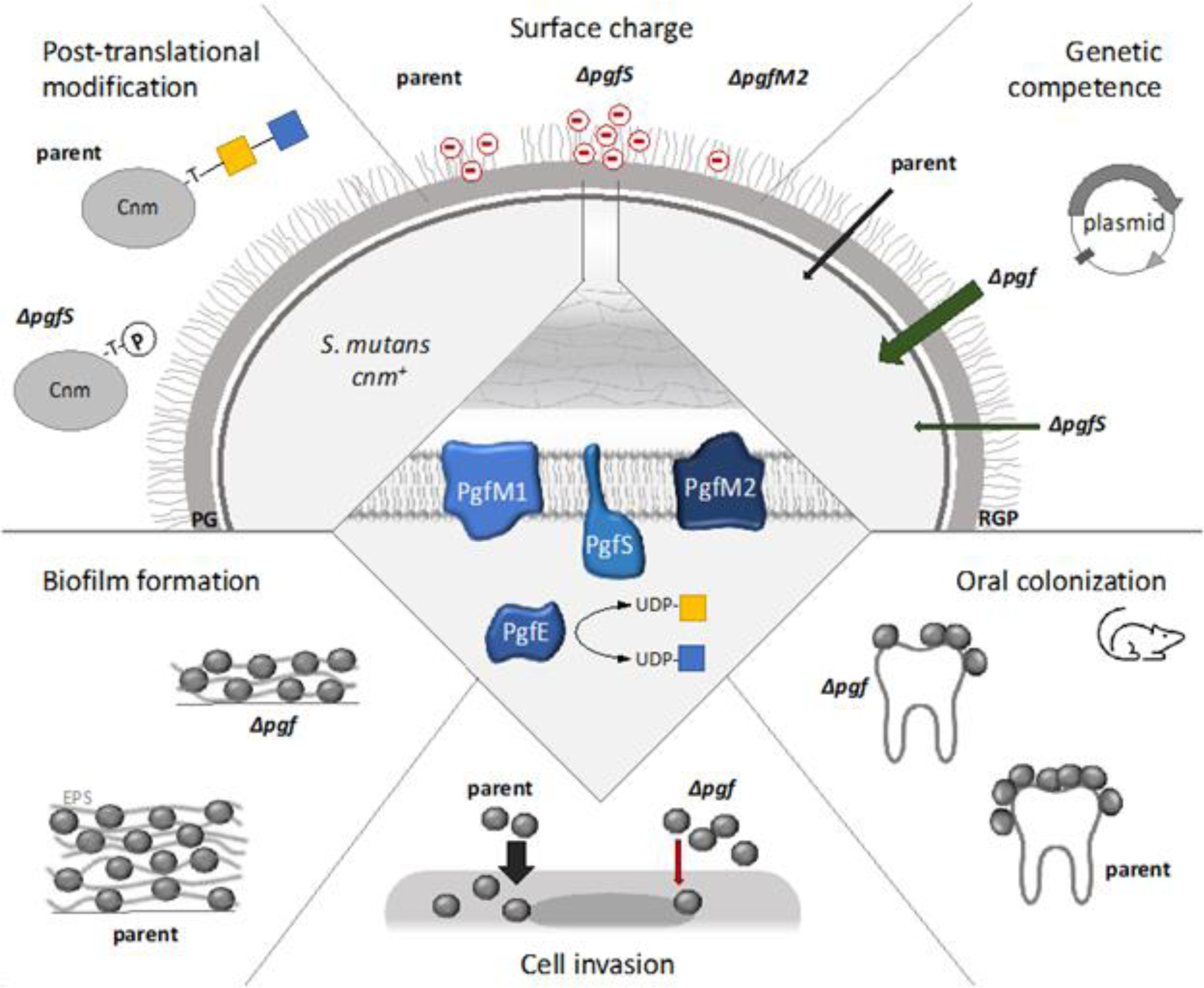

**Abbreviated summary:** In this study, we demonstrate that the Pgf glycosylation machinery of *Streptococcus mutans*, a keystone pathogen of dental caries, regulates several aspects of bacterial pathophysiology that ultimately contribute to *S. mutans* fitness in oral colonization experiments. Using the heavily glycosylated Cnm adhesin as a model, we found that inactivation of the glycosyltransferase PgfS results in loss of Cnm glycosylation, but instead, Cnm became heavily phosphorylated, suggesting a crosstalk/competition between these two post-translational modification mechanisms.

## Introduction

Living organisms require fine regulation of their biological processes by post-translational modification of proteins, which allows for rapid and often reversible modulation of protein stability and protein-substrate interactions (Prabakaran *et al*., 2012; Beltrao *et al*., 2013). Phosphorylation, glycosylation, ubiquitination, acetylation, lipidation, and many other protein modifications add useful diversity to living organisms’ toolkits (Beltrao *et al*., 2013). Glycosylation, for example, modulates protein structure, solubility, and stability, and therefore influences a myriad of cellular processes. Glycosylation can also mediate interactions with other cells and with the extracellular matrix (Spiro, 2002; Nothaft and Szymanski, 2010; Lu *et al*., 2015; Schäffer and Messner, 2017; Eichler, 2019). Not surprisingly, glycosylation is a critical post-translational modification that modulates bacterial traits associated with pathogenesis such as adhesion, motility, glycosyltransferase-based toxicity (toxic glycosylation of host proteins), host-pathogen interaction, and evasion of the immune system (Valguarnera *et al*., 2016; Bhat *et al*., 2019; Lin *et al*., 2020).

Widely present among bacteria, N- and O-glycosylations are the most common forms of protein glycosylation. The N- or -O-classification indicates the atom in the side chain of the protein that is linked to the glycan (nitrogen in asparagine and oxygen in serine and threonine). (Spiro, 2002; Nothaft and Szymanski, 2010; Nothaft and Szymanski, 2013). Both N- and O-glycosylation machineries are found in Gram-positive and Gram-negative bacteria and the addition of sugars can occur *en bloc* with the assistance of an oligosaccharyltransferase or sequentially with the assistance of glycosyltransferases (Tan *et al*., 2015). Importantly, protein glycosylation exponentially increases the complexity of the proteome, as individual proteins can be modified by oligosaccharide chains bearing different permutations of monomeric and oligomeric sugars of various sizes, charges, isomeric forms, and branching patterns (Eichler, 2019). Although these mono- and oligosaccharides are of major relevance to protein biochemistry, they are secondary gene products that are not coded in the genome, but rather covalently linked after translation by enzymes, mainly glycosyltransferases (Schäffer and Messner, 2017). This subclass of enzymes (E. C. 2.4) catalyzes the formation of glycosidic bonds between a sugar residue from a donor (most commonly a nucleotide-sugar conjugate) to an acceptor molecule such as other carbohydrates, proteins, or lipids (Schmid *et al*., 2016). There are currently 115 families of glycosyltransferases listed on CAZY (Carbohydrate-Active enZYme Database, at http://www.cazy.org) covering a wide range of biochemical processes (Lombard *et al*., 2014).

In *Streptococcus mutans*, a keystone pathogen of dental caries, glycosyltransferases (GTs), especially glycosyltransferase (Gtf) -B, -C, and -D enzymes, play a central role in pathogenicity. These GTs are part of the sucrose-dependent mechanism of colonization of *S. mutans* and are responsible for the production of the extracellular polysaccharides (EPS), a major component of the biofilm extracellular matrix (ECM). Using sucrose as a substrate, GtfB produces insoluble α(1-3)-glucans, while GtfD produces soluble α(1-6)-glucans and GtfC produces a mixture of both (Bowen and Koo, 2011; Simón-Soro and Mira, 2015; Pitts *et al*., 2017). Furthermore, Gtf enzymes can bind to microbial cell surfaces, converting other organisms into *de facto* glucan-producers, and therefore contributing to the development of thick multi-species biofilms (Krzyściak *et al*., 2014; Koo *et al*., 2018; Lemos *et al*., 2019). For instance, the binding of *S. mutans* GtfB to mannans present in the cell wall of the fungus *Candida albicans* promotes the development of a highly cariogenic biofilm (Falsetta *et al*., 2014; Hwang *et al*., 2017; Koo *et al*., 2018). Importantly, the enzymatic activity of GtfB, -C, and -D is based on the formation and breakage of glycosidic bonds between carbohydrate moieties, and not on protein glycosylation (Bowen and Koo, 2011; Krzyściak *et al*., 2014).

In addition to the aforementioned sucrose-dependent colonization mechanisms, *S. mutans* also possesses sucrose-independent mechanisms of colonization that are mainly involved in initial attachment and biofilm development (Avilés-Reyes *et al*., 2017; Arora *et al*., 2021). The surface adhesin P1, also known as Antigen I/II or SpaP, mediates *S. mutans* binding to constituents of the salivary pellicle such as the salivary agglutinin glycoprotein 340 (Gp340) and also promotes interactions between bacterial cells (Krzyściak *et al*., 2014; Sullan *et al*., 2015). The wall-associated protein A (WapA) has also been demonstrated to be important for cell-cell interaction, binding to the salivary pellicle, and organization of biofilm structure (Zhu *et al*., 2006). Both P1 and WapA were also shown to mediate collagen binding *in vitro* (Sciotti *et al*., 1997; Han *et al*., 2006). Moreover, approximately 20% of *S. mutans* clinical isolates express the collagen- and laminin-binding adhesin Cnm (Sato *et al*., 2004; Nomura *et al*., 2009; Nakano *et al*., 2010; Abranches *et al*., 2011; Avilés-Reyes *et al*., 2014b), an important virulence factor associated with dental caries severity and recurrence (Miller *et al*., 2015; Esberg *et al*., 2017; Garcia *et al*., 2021) that also contributes to extra-oral infections such as cerebral microbleeds, hemorrhagic stroke and infective endocarditis (Abranches *et al*., 2011; Nomura *et al*., 2013; Miller *et al*., 2015; Avilés-Reyes *et al*., 2017; Inenaga *et al*., 2018; Hosoki *et al*., 2020; Nomura *et al*., 2020; Arora *et al*., 2021). In addition to binding to collagen, Cnm, WapA and P1 have also been characterized as functional amyloids that help model the biofilm structure (Oli *et al*., 2012; Besingi *et al*., 2017; di Cologna *et al*., 2021; Yarmola *et al*., 2022). In addition to sharing functional and structural similarities, Cnm and WapA were both shown to be post-translationally modified through glycosylation (Avilés-Reyes *et al*., 2017; Avilés-Reyes *et al*., 2018).

Post-translational modifications are increasingly implicated as having relevant functional roles in the pathogenesis of oral bacteria (Ma *et al*., 2021). In *S. mutans* OMZ175, the *cnm* gene is co-transcribed with the four-gene *pgf* operon (*pgfS*, *pgfM1*, *pgfE,* and *pgfM2*) that is part of the *S. mutans* core genome and is required to glycosylate both core (WapA) and non-core (Cnm) LPXTG-anchored surface proteins (Avilés-Reyes *et al*., 2018). Like *cnm,* the *pgf* operon is negatively regulated by VicRKS and positively regulated by CovR, two signal transduction systems that control the expression of many virulence-associated genes of *S. mutans* (Stipp *et al*., 2013; Avilés-Reyes *et al*., 2018; Alves *et al*., 2018). In *S. mutans* OMZ175, deletion of the *pgf* genes, either individually or the entire operon, resulted in a reduction of the observed mass of Cnm and WapA in polyacrylamide gels, and both proteins demonstrated reduced stability and resistance to proteolysis. Also, deletion of the *pgf* genes led to a significant decrease in expression of Cnm-associated virulence traits including collagen-binding, invasion of endothelial and epithelial cells, and killing of the invertebrate *Galleria mellonella* model of systemic infection (Avilés-Reyes *et al*., 2014a; Avilés-Reyes *et al*., 2018).

To gain further insights into post-translational glycosylation mechanisms employed by the Pgf machinery and the contribution of each Pgf protein to *S. mutans* fitness, we performed a series of phenotypic assays to study important traits associated with *S. mutans* pathophysiology using a panel of *pgf* mutants created in the *cnm*^+^ strain OMZ175. Additionally, using the glycoprotein Cnm as a Pgf substrate model, we performed glycoproteomic analyses of a series of Cnm constructs expressed in OMZ175 and the *pgf* mutants.

## Results and Discussion

### The Pgf glycosylation system is responsible for many aspects of bacterial pathobiology

Our previous studies focused on the role of the Pgf glycosylation system on the expression of Cnm-dependent phenotypes such as collagen- and laminin-binding and intracellular invasion (Avilés-Reyes et al., 2014a; Avilés-Reyes et al., 2018). However, considering that Pgf is also present in *cnm*-negative strains and that it modifies the core adhesin WapA, we anticipated that the scope of the Pgf system goes beyond Cnm and WapA modification (Avilés-Reyes *et al*., 2014a; Avilés-Reyes *et al*., 2018). Current and past phenotypical characterization of mutants for the Pgf machinery revealed a wide spectrum of changes in phenotype as compared to the parental strain and complemented strains (Avilés-Reyes *et al*., 2014a; Avilés-Reyes *et al*., 2018), summarized in Table 1.

**Table 1.**
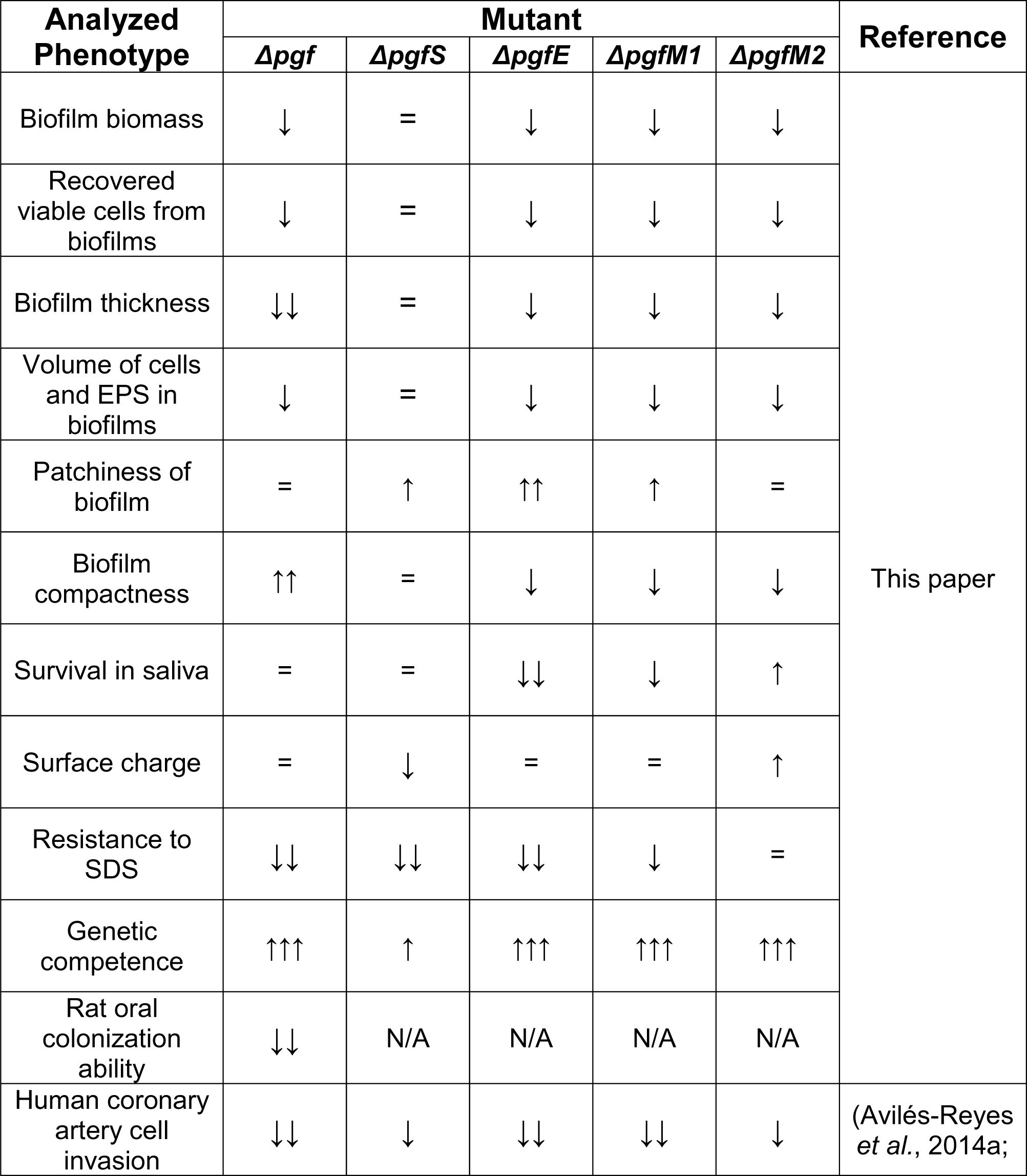
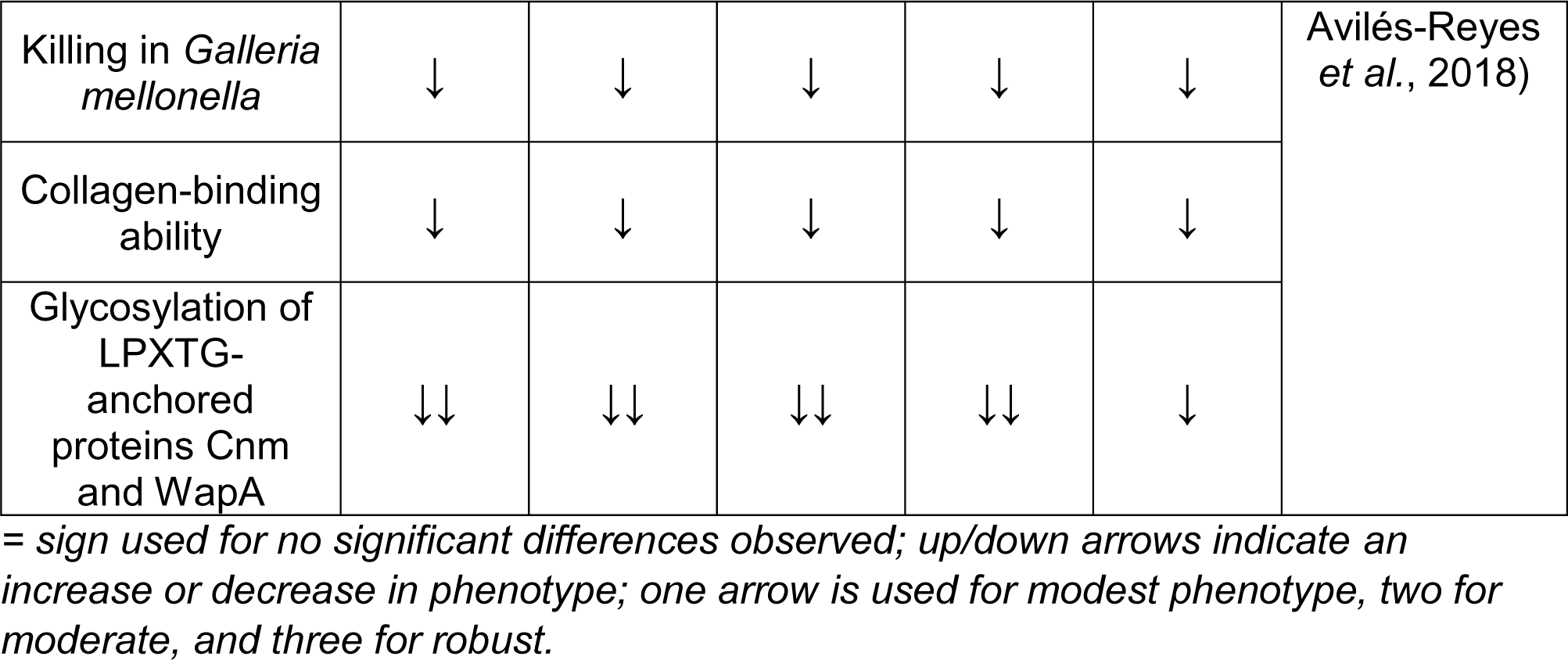
Summary of phenotypical differences observed for the pgf mutants as compared to the parental strain OMZ175 in current and previous studies.

Carbohydrates are essential for *S. mutans* pathobiology, as this organism relies mostly on sugars for energy generation, biofilm formation, and building the cell wall and its associated polysaccharides such as the rhamnose-glucose polysaccharide (RGP) layer (Abranches *et al*., 2018; Lemos *et al*., 2019). There are several predicted glycosyltransferases in the genome of *S. mutans* including those from the RGP synthesis pathway (RgpA, RgpB, RgpE, RgpF, RgpH, and RgpI) (Shibata *et al*., 2009; Mistou *et al*., 2016; Rainey *et al*., 2018; Kovacs *et al*., 2019; Bischer *et al*., 2020). Other uncharacterized putative glycosyltransferases are annotated in the *S. mutans’* genome including Smu.834, located immediately downstream of the *rgp* operon (Lara Vasquez *et al*., 2021), Smu1039c, Smu1434c, and Smu1806. Therefore, the Pgf proteins may not be the only designated protein glycosylation machinery in *S. mutans*, raising the possibility of functional redundancy between pathways and highlighting the importance of these systems for bacterial pathobiology.

### *In silico* analyses of the Pgf proteins offer hints related to their structure and function

Our first step in understanding the function of the Pgf machinery was by expanding the bioinformatics analysis initiated in our previous study (Avilés-Reyes *et al*., 2018). In addition to domain prediction using InterPro (Blum *et al*., 2021) and domain localization of transmembrane proteins using TMHMM 2.0 (Krogh *et al*., 2001), the current state-of-the-art predictor of protein structure AlphaFold2 (Jumper *et al*., 2021) was used in HiPerGator 3.0, the University of Florida’s supercomputer. AlphaFold2 has been demonstrated as the best currently available tool for the prediction of the 3D structure of proteins for which no homologous structure is known (the “protein folding problem”) in the latest Critical Assessment of protein Structure Prediction (CASP, 14^th^ edition), and it operates utilizing a neural network-based machine learning algorithm that interprets evolutionary, biophysical, and steric constraints starting from a protein sequence input (Jumper *et al*., 2021). Together, these analyses indicate that PgfS, PgfM1, and PgfM2 are likely membrane proteins, while PgfE is an intracellular globular protein (Figure 1A). Moreover, PgfS contains an intracellular nucleotide-diphospho-sugar-transferase domain, while PgfM1 and PgfM2 have an extracellular globular domain of no predicted function. PgfE is predicted to contain a Rossmann Fold, a highly conserved motif consisting of up to seven strands of alternating β-sheets and α-helixes which interact via hydrogen bonds (Hanukoglu, 2015). This domain is well known for binding NAD(P), and FAD dinucleotides (Hanukoglu, 2015), which suggests the possibility of these dinucleotides being coenzymes of PgfE. GT-A nucleotide-sugar-dependent glycosyltransferases such as PgfS (Avilés-Reyes *et al*., 2014a) and the GalT proteins of oral streptococci are known to contain a single Rossmann fold (Zhu *et al*., 2015), and the presence of a Rossmann Fold motif is suggested by AlphaFold2 modeling of PgfS. Initial analysis by InterPro revealed that the homologous membrane proteins PgfM1 and PgfM2 (88% coverage, with 38.5% identity and 54% positives when aligned) are not predicted to contain a Rossmann fold. However, AlphaFold2 structure predictions of these proteins also display alternating β-sheets and α-helixes in their extracellular globular domains (Figure 1A), suggesting that the entire Pgf machinery could be relying on NAD(P) and/or FADH as coenzymes. An *in silico* homo-oligomerization analysis using the AlphaFold2 predicted structures of Pgf proteins in the GalaxyHomomer platform (Baek *et al*., 2017) predicts that PgfS likely forms a tetramer, while PgfE likely forms a dimer (Figure 1B). The template-based prediction of the PgfS tetramer was based on the GtrB polyisoprenyl-phosphate glycosyltransferase from *Synechocystis* sp recombinantly expressed in *Escherichia coli* (PDB ID 5EKP), with an interface area of 7662.2 Å^2^ between monomers. GtrB and PgfS share a 34.7% sequence identity, and a 76.2% structural similarity. The crystal structure of the GtrB transmembrane tetramer was previously determined via X-ray crystallography with 3.19 Å of resolution (Ardiccioni *et al*., 2016). The dimer prediction of PgfE was based on the UDP-4-galactose epimerase GalE structure from *Homo sapiens* (PDB ID 1I3K), an enzyme of the Leloir pathway for galactose catabolism that also exists in *S. mutans*, with 920.3 Å^2^ of interface area between monomers. The PgfE and the human GalE proteins share 53.6% of sequence identity and 98.26% of structural similarity. The crystal structure of the human GalE was also determined via X-ray crystallography with 1.50 Å of resolution (Thoden *et al*., 2001). A GalaxyHomomer analysis of the AlphaFold2 structures from PgfM1 and PgfM2 did not return any expected oligomers.

**Figure 1.**
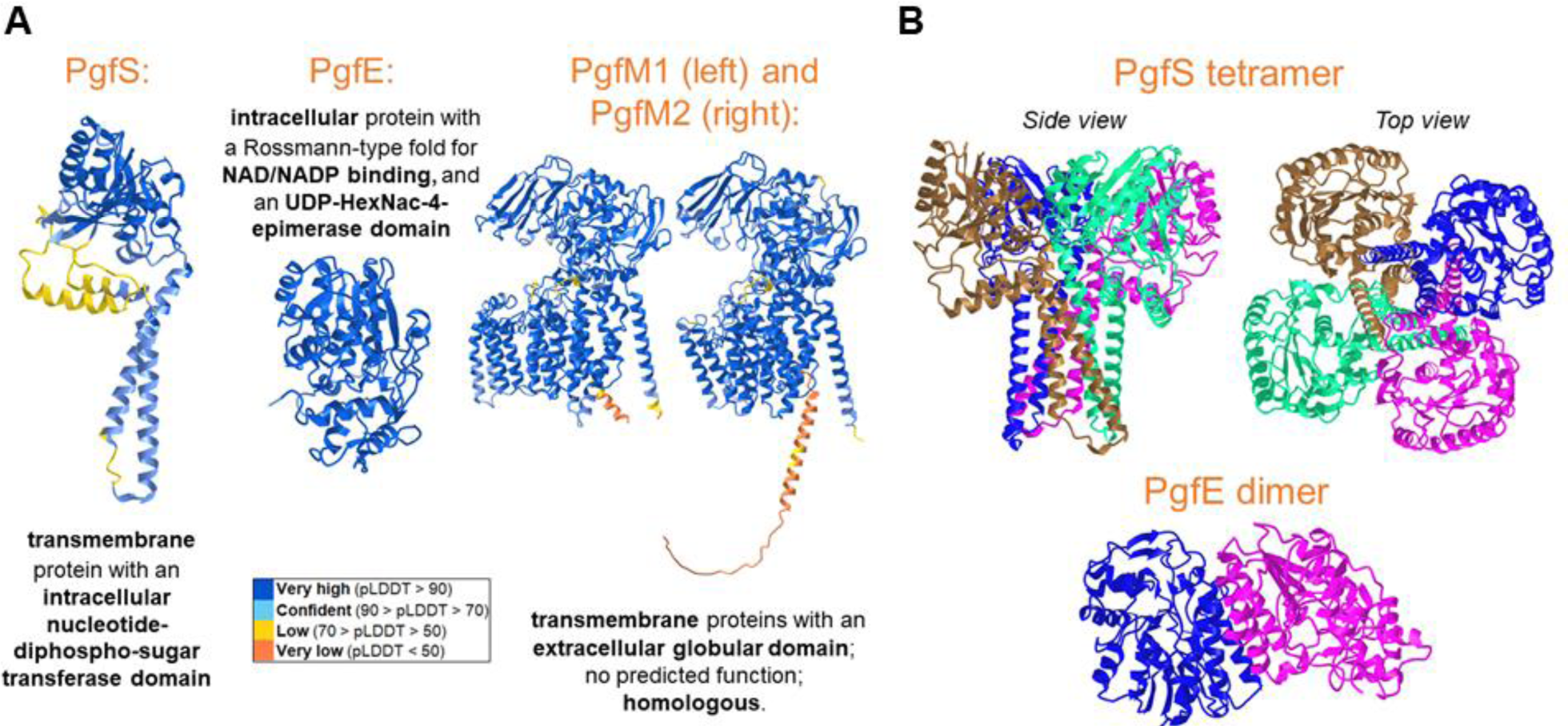
Bioinformatics prediction for structural features of Pgf proteins. (A) Monomer analysis via Alpha Fold (structure prediction), InterPro (Domains), and TMHMM 2.0 (prediction of domain positions in transmembrane proteins). Colors indicate AlphaFold per-residue estimate of confidence (pLDDT). (B) GalaxyHomomer template-based modeling for Pgf proteins predicts PgfS as a tetramer (template PDB - 5EKP) and PgfE as a dimer (template PDB - 1I3K). Each color indicates a different monomer.

### Glycosylation by the Pgf machinery is required for optimal biofilm formation and possible crosstalk with other glycosylation systems

To gain insight into the importance of the Pgf machinery in *S. mutans’* key virulence traits, we analyzed the biofilm formation ability of our panel of mutant strains (*ΔpgfS, ΔpgfE, ΔpgfM1, or ΔpgfM2* strains and *Δpgf* which designates a strain with all four *pgf* genes deleted). Figure S1 displays a schematic diagram of the mutant strains utilized in our experiments and of the *cnm-pgf* locus in OMZ175. When biofilms were allowed to grow for 48 hours in a chemically defined biofilm medium (CDM) supplemented with 1% sucrose, all of the mutant strains showed a defect in biofilm biomass when compared to the parent strain, except for *ΔpgfS* (Figure 2A). Previously, we showed that the inactivation of *cnm* in OMZ175 resulted in decreased biofilm formation in media containing sucrose (Abranches *et al*., 2011). Thus, the observed phenotype for the mutants could be, at least in part, related to Cnm function and stability, since this adhesin is a known target of the Pgf glycosylation machinery. However, the *ΔpgfS* mutant phenocopied the parent strain in the biofilm biomass assay, and because PgfS is required for full Cnm glycosylation (Avilés-Reyes *et al*., 2018), it is likely that the lack of proper protein glycosylation also affects biofilm biomass in a Cnm-independent manner. A similar trend was observed for the recovery of viable cells from these biofilms (Figure 2B), with comparable CFUs recovered from the parent strain and *ΔpgfS* mutant biofilms and significantly lower cells recovered from the other *pgf* mutant biofilms. Thus, our findings indicate that proper protein glycosylation is important for the ability of *S. mutans* to form robust sucrose-dependent biofilms.

**Figure 2.**
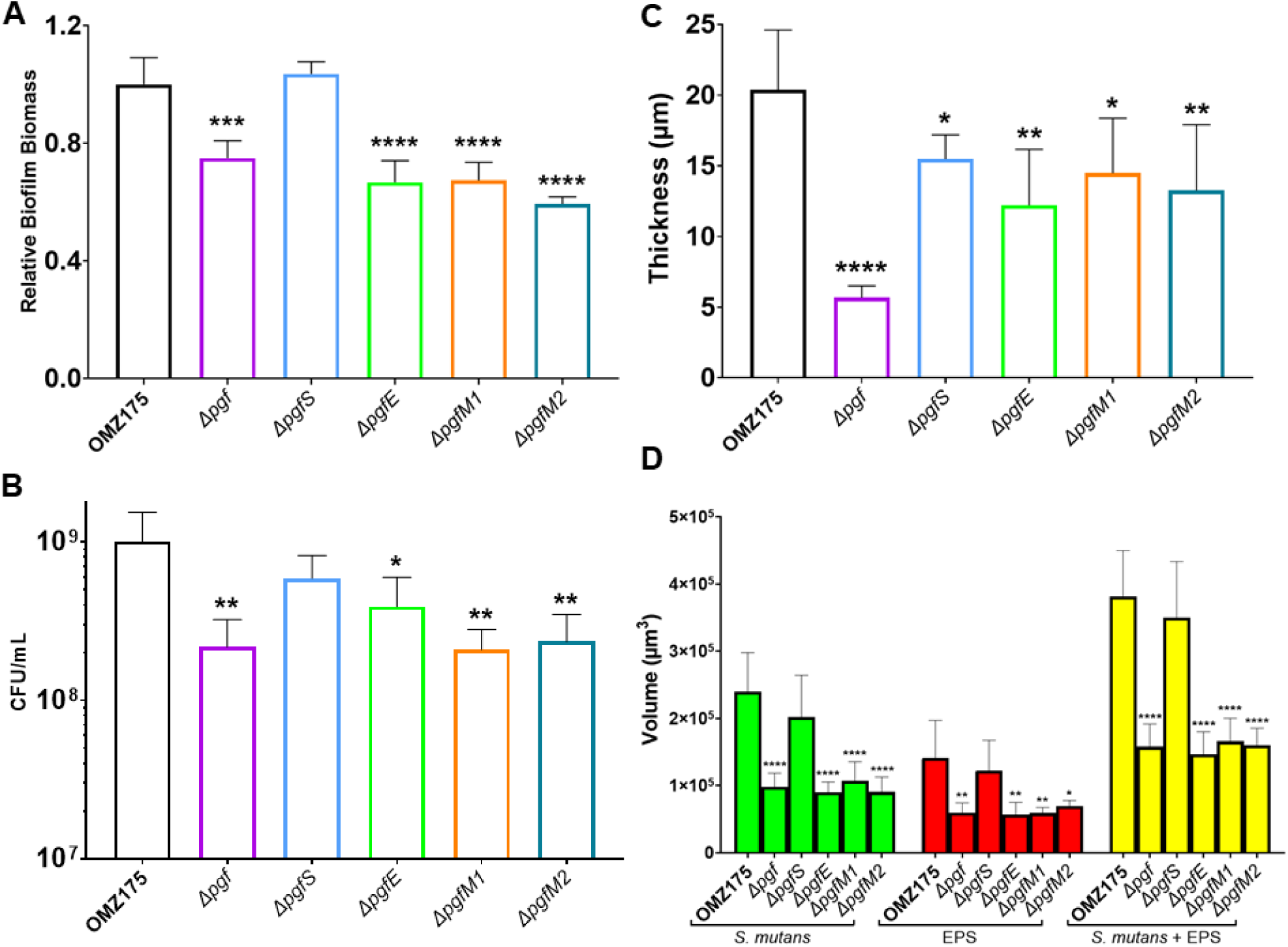
The Pgf glycosylation machinery is important for proper biofilm formation. Deletion of the entire *pgf* operon or each gene individually, except for *pgfS*, results in biofilm biomass defects. (A) Crystal violet staining of biofilms relative to the parental strain, measured by absorbance at 595 nm. (B) Viable cells recovered from biofilms. (C) Biofilm thickness as measured on a volumetric view via Confocal Laser Microscopy Scanning (p < .001). (D) The estimated volume of biofilms was calculated from Confocal Laser Microscopy Scanning by multiplying each area by the distance between layers. (p < .0001 for *S. mutans*; p = .0003 for EPS; p < .0001 for sum). Each group was analyzed via one-way ANOVA. Asterisks denote post-hoc comparison with the parental strain. * = p < .05; ** = p < .01; *** = p < .001; **** = p < .0001. Bars indicate mean values and error bars indicate standard deviations. N = 4.

The architecture of *S. mutans* biofilms is largely dependent upon the production of a matrix whose components act as scaffolds that guide the three-dimensional distribution of microenvironments within the biofilm (Klein *et al*., 2015). To further dissect the effects of the *pgf* deletions in biofilm formation, confocal laser microscopy scanning (CLMS) of all strains was performed using labeling fluorophores for *S. mutans* cells and extracellular polysaccharides. To quantitatively analyze the characteristic parameters of these biofilms, each z-stack was converted into a binary layer and the data were used for an in-depth quantitative analysis of the biofilms (Figure 2C, D, and Figure S2). When compared to the parent strain, the overall biofilm architecture was affected in each *pgf* mutant tested (Figure 3). Although all quantitative metrics for the comparison between the parent strain and the *ΔpgfS* mutant were comparable as indicated above, when the complex 3D structure is visualized, it is noticeable that the parent strain’s biofilm is more homogeneously distributed over the area in each layer, while some degree of patchiness and larger microcolonies are observed for *ΔpgfS, ΔpgfE,* and *ΔpgfM1*. The *ΔpgfM2* biofilm, however, was homogeneously distributed, similar to the parent strain. Finally, the *Δpgf* quadruple mutant showed a thinner, more compact, and more homogeneous biofilm when compared to the parent and other mutant strains. Changes in biofilm thickness (distance between bottom and top of biofilms) and volume of components (cells and EPS) were the most striking phenotypes observed for these mutants. The parent strain biofilm was thicker than all mutants’ biofilms except for *ΔpgfS*, and the *Δpgf* mutant biofilm appears to be the thinnest of the biofilms as seen in Figure 2C. The volume of the biofilms (cells, EPS, and sum) was also estimated (Figure 2D). All mutants presented an overall lower volume of cells, EPS, and their sum than the parental strain, except for the *ΔpgfS* mutant. Thus, all *pgf* genes are important for robust biofilm development whereas *pgfS* appears to contribute to a lesser extent. Despite the differences in architecture, the volumetric proportion of each analyzed component remained at approximately a 3:2 ratio (*S. mutans*:EPS) for all analyzed biofilms (Figure S2A). Lastly, biofilm compactness was estimated, indicating high compactness of the *Δpgf* mutant (Figure S2B). The observation that the *ΔpgfS* mutant phenocopies the parental strain is intriguing, considering that it is the only member of this glycosylation machinery with a traditional GT domain. This observation raises the possibility of functional redundancy of PgfS with other *S. mutans* glycosyltransferases. A protein BLAST search against an *S. mutans* database using the PgfS amino acid sequence revealed only two significant hits: glucosyltransferase RgpI (95% coverage with 41% identities and 61% positives) and rhamnosyltransferase RgpB (36% coverage with 28% identities and 47% positives), both enzymes from the RGP synthesis machinery (Kovacs *et al*., 2019). The RGP-derived modifications of the *S. mutans* cell wall are important in guaranteeing proper septation and location of cell division complexes, and also in preventing autolysis (Kovacs *et al*., 2019; Zamakhaeva *et al*., 2021). RgpI has been demonstrated as having a role in modulating the matrix of biofilms in *S. mutans*, mainly by decreasing glucan components and increasing the abundance of extracellular DNA (eDNA), without altering the overall biomass (Rainey *et al*., 2018). Thus, a similar effect may have occurred for the *ΔpgfS* mutant, where a change in biofilm architecture, but not biofilm biomass is observed (Figures 2 and 3). The AlphaFold2 predicted structure for RgpI is also very similar to that of PgfS (Figure S5). Therefore, we cannot rule out that PgfS and RgpI may have overlapping roles in promoting appropriate biofilm development. Taken altogether, our findings show an important role of the Pgf glycosylation machinery in biofilm accumulation and architecture, and a possible functional redundancy between PgfS and RgpI as further discussed below.

**Figure 3.**
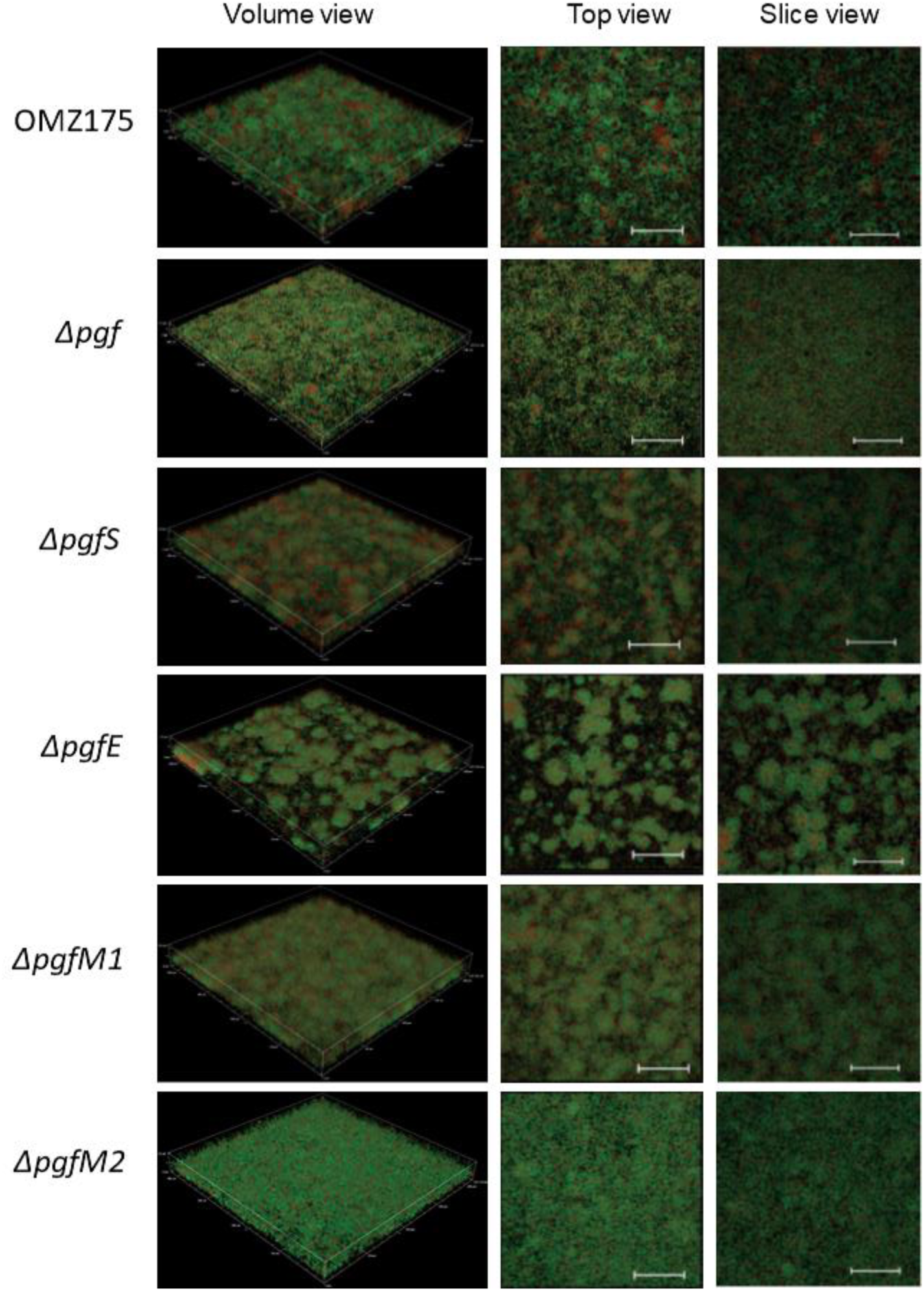
A proper glycosylation state is important for biofilm assembly and architecture. Confocal Laser Scanning Microscopy of the panel of *S. mutans* strains in volumetric view displays architectural differences for all *pgf* mutants when compared to the parental strain OMZ175. A top (from above) and a slice view (middle of biofilm) are also represented. Representative images were chosen from four biological replicates, with three images collected for each. Green = *S. mutans* cells, Red = Extracellular Polysaccharides (EPS). Each side of the square base = 210 µm. Scale bar = 50 µm.

### Adequate glycosylation by the Pgf (and possibly RGP) system interferes with *S. mutans* survival in human saliva, surface charge, and membrane stability

In order to establish a biofilm in the oral cavity, microorganisms must be able to utilize salivary components as nutrients as well as tolerate salivary secretory immunity, comprised of soluble immunoglobulins (sIgA and, to a lesser extent, sIgM), proteases, and salivary antimicrobial proteins and peptides (Amorim *et al*., 2011; Fábián *et al*., 2012; Brandtzaeg, 2013; Lynge Pedersen and Belstrøm, 2019). When we measured the survival in saliva (supplemented with glucose to prevent starvation-related cell death) of our panel of strains and compared it to that of the parent strain, we observed that the absence of Pgf proteins, except for PgfM2, significantly diminished survival in saliva (Figure 4). Despite the high homology between PgfM1 and PgfM2, the *ΔpgfM1* strain was almost entirely cleared after 20 hours of incubation (≈0.1% survival, our limit of detection), whereas the *ΔpgfM2* mutant only crossed the 0.1% survival threshold sometime between 28-48 hours of incubation. Interestingly, the *ΔpgfE* mutant was the most susceptible and was below the detection limit after 20 hours of incubation, indicating an important role of the regulation of the availability of GlcNAc and GalNAc for the bacterial ability to survive in saliva. Also, the quadruple Δ*pgf* mutant showed a significant reduction in survival after 28 hours of incubation in saliva. Post-translational modification of bacterial surface proteins by phosphorylation, lipidation, glycosylation, and others is used by bacteria to evade the immune system. For example, bacteria can be cloaked in hydrophobic, hydrophilic, or charged molecules that repel antimicrobial peptides and proteins, while molecular mimicry of human protein-associated glycans avoids recognition by antibodies (Reddick and Alto, 2014; Brockhausen, 2014; Macek *et al*., 2019). Since at least two of the known targets of the Pgf machinery are surface-anchored proteins, it is not surprising that the mutants behave differently than the parent strain when in the presence of saliva. Deletion of PgfM2 still results in partially glycosylated Cnm and WapA, while the deletion of any other Pgf protein resulted in fully unglycosylated Cnm and WapA (Avilés-Reyes *et al*., 2018), and this may be the reason why the survival in saliva of the *ΔpgfM2* strain was not compromised as it was for the other mutants. UDP-4-galactose epimerases have been demonstrated as crucial agents in cell surface glycocomponent generation in both Prokaryotes and Eukaryotes (Fry *et al*., 2000; Roper *et al*., 2005; Lee *et al*., 2014; Schäper *et al*., 2019; Broussard *et al*., 2020). Thus, exposed cell envelope-linked glycans are likely important for the recognition of this bacteria by salivary antimicrobial components.

**Figure 4.**
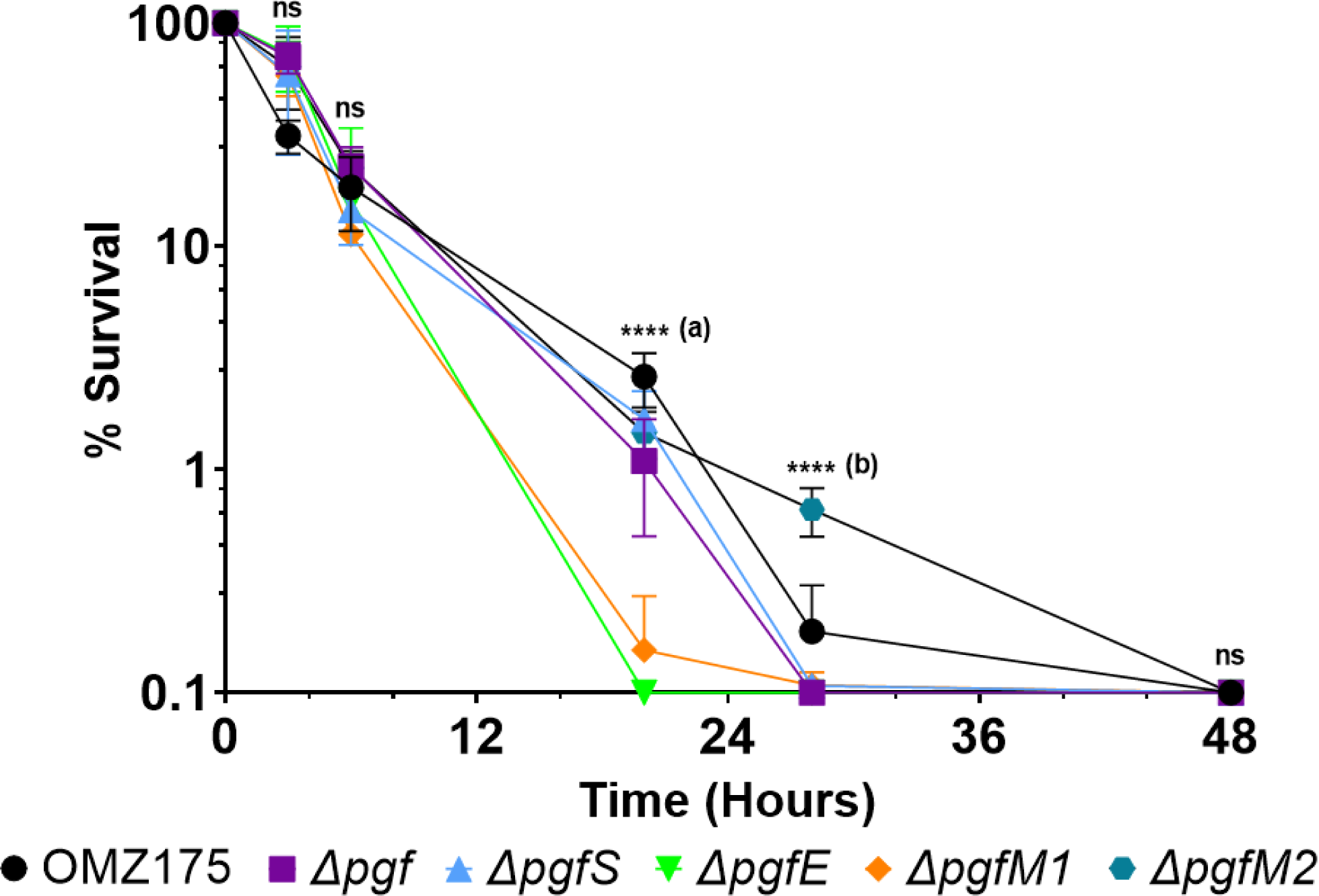
The Pgf glycosylation machinery contributes to *S. mutans* survival in human saliva supplemented with 20 µM of glucose. The deletion of any *pgf* gene or all of them decreased the survival of bacteria in human saliva, except for the deletion of only *pgfM2*. 0.1% survival was the limit of detection. Connected dots indicate means, and error bars indicate standard deviations. One-way ANOVA was performed on each timepoint to determine differences between each mutant and the parental strain. Asterisks denote post-hoc comparison with the parental strain. (a) denotes differences between parental and *ΔpgfS/ΔpgfM2* (p < .05) and between parental and *Δpgf/ΔpgfE/ΔpgfM1* (p < .0001). (b) denotes differences between parental and ΔpgfM2 only (p < .0001). N = 4. ns = non-significant; * = p < .05; ** = p < .01; *** = p < .001; **** = p < .0001.

We next directed our efforts to the characterization of cell envelope homeostasis in the *pgf* mutants. First, we estimated the cell surface charge via zeta potential measurement (Figure 5A), which serves as an indirect measurement of the electrical potential between the bacterial surface and the surrounding water pellicle (Wilson *et al*., 2001). Of note, proper surface charge is pivotal to the maintenance of a surface hydrophobicity that allows for transmembrane transport of nutrients and waste in an aqueous environment (Wilson *et al*., 2001). Gram-positive lactic acid bacteria present negatively charged surfaces under physiological pH due to the presence of lipoteichoic acids (LTAs) and wall teichoic acids (WTAs) imbued within the peptidoglycan layer (Wilson *et al*., 2001; Chapot-Chartier and Kulakauskas, 2014). In *S. mutans*, there are no WTAs and the RGP layer is considered to fulfill their roles, although it does not contain the negative charge associated with WTA (Kovacs *et al*., 2019; Bischer *et al*., 2020). When compared to the parental strain, the deletion of *pgfS* significantly increased the negative charge by ≈44%, while the deletion of *pgfM2* significantly decreased the negative charge by ≈26%. On the other hand, the deletion of *pgfE* and *pgfM1* did not significantly change the bacterial zeta potential. Of note, the deletion of the entire *pgf* operon (*Δpgf*) did not alter the cell surface charge. Therefore, the simultaneous deletion of *pgfS* and *pgfM2* in *Δpgf* appears to erase the opposing effects of each individual mutation. We hypothesized that this surface charge variation was due to a modification of the glycosylation pattern of surface proteins or, perhaps, other constituents of the cell envelope like lipids and the RGP. In addition to PgfS being nearly identical to the glycosyltransferase RgpI of the RGP synthesis pathway (Figure S5), the PgfS and PgfM1/M2 proteins are also highly homologous at both structural and sequence levels to LTA and WTA glycosylation enzymes characterized in other Gram-positive bacteria, such as *Listeria monocytogenes* (GtlA/Lmo2550 as PgfS-like and GtlB/Lmo1079 as PgfM1/M2-like proteins, respectively) and *Bacillus subtilis* (CsbB and YfhO as PgfS-like and PgfM1/M2-like proteins, respectively) (Rismondo *et al*., 2018; Rismondo *et al*., 2020; Rismondo *et al*., 2021). Figure S6 depicts and compares the predicted structures for PgfS and GtlA, Lmo2550 and CsbB, and Figure S7 depicts the same comparison for PgfM1, PgfM2, Lmo1079, GtlB, and YfhO. Also, as previously shown in Figure 1B, the PgfS predicted tetramer is structurally similar to the GtrB tetramer from *Synechocystis* sp., a well-characterized and conserved polyisoprenyl-phosphate glycosyltransferase (PGT) (Ardiccioni *et al*., 2016). PGTs are integral membrane proteins that catalyze the transfer of a sugar moiety from a nucleotide-sugar to undecaprenyl-phosphate (in Bacteria) or dolichol-phosphate (Eukarya and Archaea), which act as transmembrane carriers of sugar to allow for the formation of membrane glycoconjugates (Ardiccioni *et al*., 2016; Allen and Imperiali, 2019). The surface charge of teichoic acids can be modulated by replacing free hydroxyl groups of their alditol-phosphate with carbohydrates or D-alanine residues (Chapot-Chartier and Kulakauskas, 2014). The presence or absence of neutral or charged carbohydrates in the membrane, as well as the relative abundance of exposed peptidoglycan/LTA (negatively charged) when compared to the glycosylated peptidoglycan/LTA, may alter the cell’s zeta potential and overall membrane stability. Further research is required to determine whether the Pgf machinery can also glycosylate membrane lipids since the speculations made here are solely based on *in silico* analysis. Importantly, the characterized LTA/WTA glycosylation machineries do not possess a designated sugar epimerase like PgfE, but, surprisingly, PgfE is 38.85% identical (56% positives, 99% coverage) to the glucose-galactose epimerase GalE of the Leloir pathway in *S. mutans*. Our group recently characterized and compared the functions of PgfE and GalE and found that despite their structural similarities, PgfE mostly recognizes N-acetylated sugars as substrates (GlcNAc and GalNAc), while GalE only recognizes non-acetylated sugars (Glc and Gal) (Andresen *et al*., 2022a). When we tested our panel of strains for resistance to detergents, significant differences in phenotypes were seen only for the negatively charged detergent sodium dodecyl sulfate (SDS). Interestingly, the Δ*pgfS*, Δ*pgfE,* and the quadruple Δ*pgf* mutant presented a significantly increased sensitivity to SDS as measured by minimum inhibitory concentration (MIC) determination (Figure 5B, top and Figure S3) whereas the *ΔpgfM1* and *ΔpgfM2* mutants phenocopied the parent strain. We speculate that the fact that the *ΔpgfM2* mutant (and possibly *ΔpgfM1* as well) surface is less negatively charged than the parental strain’s, this difference in surface charge allows for a partial neutralization of the disturbance that SDS inflicts upon the cellular membrane. No differences in MIC were observed for non-ionic detergents Tween-20 and Triton X-100 (data not shown). To test the hypothesis that the *pgf* operon is also acting on RGP glycosylation, we utilized the RGP/peptidoglycan synthesis inhibitor tunicamycin (Zhu *et al*., 2018; Kovacs *et al*., 2019) for MIC determination (Figure 5B, bottom and Figure S4). The high sensitivity of most *pgf* mutants, especially the quadruple mutant *Δpgf*, is suggestive of the participation of the Pgf machinery in RGP synthesis. Thus, our findings may indicate that the scope of the Pgf machinery might go beyond glycosylation of surface adhesins, also contributing to cell wall synthesis and membrane homeostasis. Future studies including an *ΔrgpI* mutant and a double *ΔpgfSΔrgpI* mutant could help us understand how redundant those two enzymes can be, and perhaps explain why the *ΔpgfS* biofilms were similar to the parental strain biofilms. Other phenotypic traits of interest including antibiotic disc inhibition, ethidium bromide permeability, survival in serum, and opsonophagocytosis susceptibility were analyzed, but all mutants phenocopied the parental strain (Figure S8).

**Figure 5.**
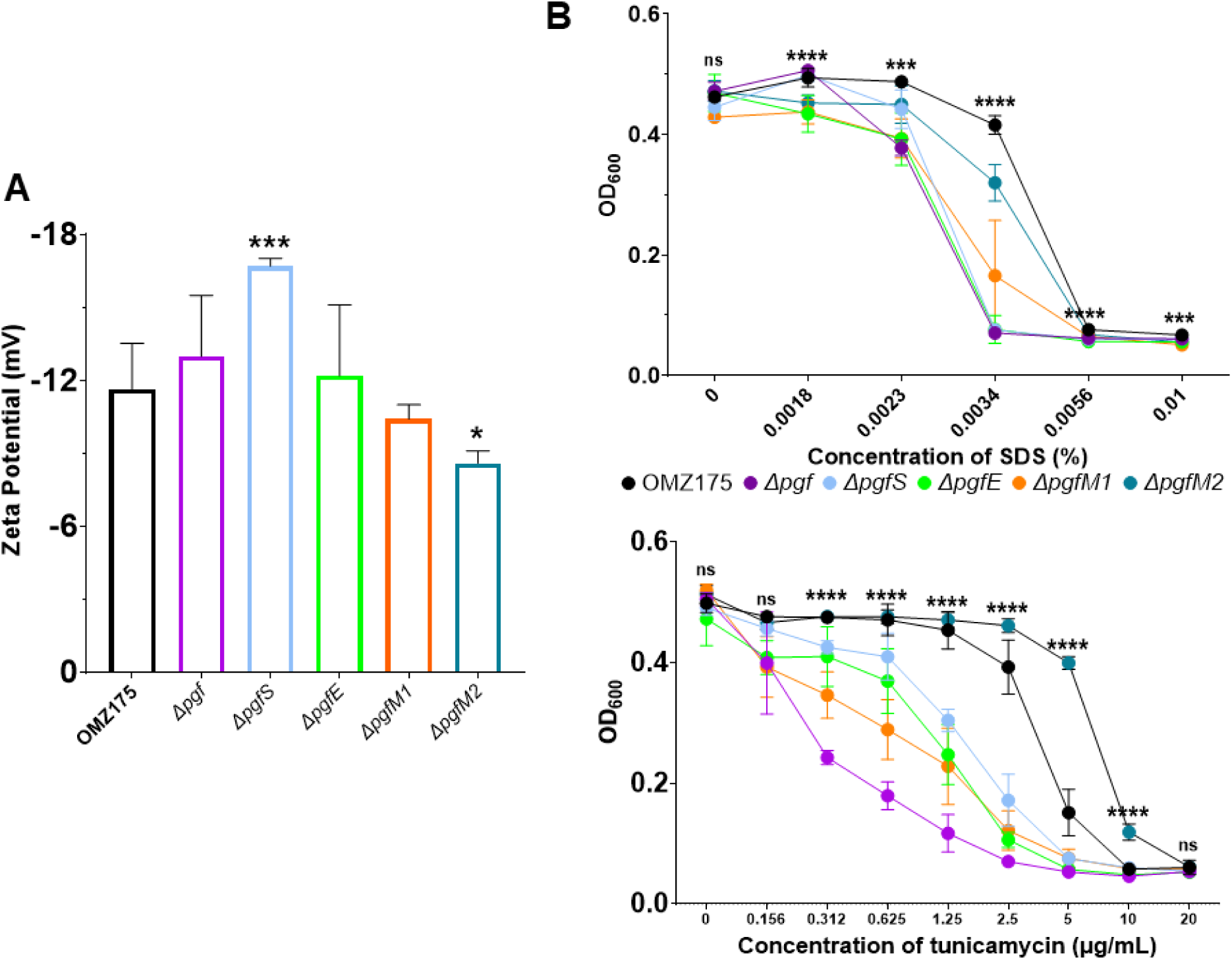
The Pgf machinery contributes to bacterial surface charge and stability. (A) Zeta potential of cells grown to mid-log phase. (B) MIC determination for the negatively charged detergent SDS (top) and the rhamnose-glucose polysaccharide synthesis inhibitor tunicamycin (bottom). Bars and dots indicate mean values, and error bars indicate standard deviations. One-way ANOVA was performed on each timepoint to determine differences between each mutant and the parental strain. Asterisks denote post-hoc comparison with the parental strain. Pairwise comparisons for growth in SDS and tunicamycin can be found in Figure S3 and Figure S4, respectively. N = 4. ns = non-significant; * = p < .05; ** = p < .01; *** = p < .001; **** = p < .0001.

### Transformation ability via the ComABCDE and ComRS pathways is suppressed by the Pgf machinery

The phenotypes observed for membrane homeostasis prompted us to also analyze genetic competence, a process dependent on the ability of surface-associated proteins to bind and take up DNA (Dubnau and Blokesch, 2019). The ability to share genetic material through competence is a keystone event in streptococcal pathogenesis (Sitkiewicz, 2018; Kaspar and Walker, 2019). *S. mutans* is a naturally competent bacterium, with the competent state triggered by activation of the alternative sigma factor ComX via the interconnected ComABCDE and ComRS pathways (Son *et al*., 2015). These pathways are activated by the competence-stimulating peptide (CSP, encoded by *comC*) and by the *comX*-inducing peptide (XIP, encoded by *comS*) in response to stresses and environmental conditions (Son *et al*., 2015; Underhill *et al*., 2018; Underhill *et al*., 2019). The regulation of genetic competence in *S. mutans* is complex and there are still gaps in knowledge. Transformation assays were performed using CSP or XIP to assess whether the Pgf glycosylation machinery can influence these competence pathways in *S. mutans* (Figure 6). Our findings revealed that when compared to the parent strain, a 3-log (CSP) and 2-log (XIP) increase in transformability for all mutants, except for *ΔpgfS*, was observed, suggesting that the optimal glycosylation state conferred by the Pgf machinery dampens competence. Genome-wide screening of a transposon library in *S. mutans* identified several genes that, when deleted, induce an aberrant expression of *comX* (Shields *et al*., 2018). In addition to the known competence-related genes, other genes involved in cell division and cell envelope homeostasis were identified (resistance to stresses and biogenesis), suggesting that they are likely important for the activation of early competence (Shields *et al*., 2018). Interestingly, *rgpI* was among the genes identified in the transposon library with increased competence, which due to its high homology with *pgfS,* raises the possibility of the latter also influencing DNA uptake. However, our data revealed only a modest, non-significant increase in competence for the *ΔpgfS* mutant, once again suggesting a possible functional redundancy between PgfS and RgpI in the expression of some bacterial traits. Moreover, RGP glycosylation has been recently shown to be a crucial step for the recruitment of the cell-division machinery (Zamakhaeva *et al*., 2021) and cell growth (Bischer *et al*., 2020), strengthening the link between protein glycosylation, cell division, and genetic competence. An alternative explanation for the small increase in transformability observed for the *ΔpgfS* mutant when compared to other mutants is that the more negative surface charge of this mutant (Figure 5A) could be repelling the negatively charged DNA, inhibiting its intake.

**Figure 6.**
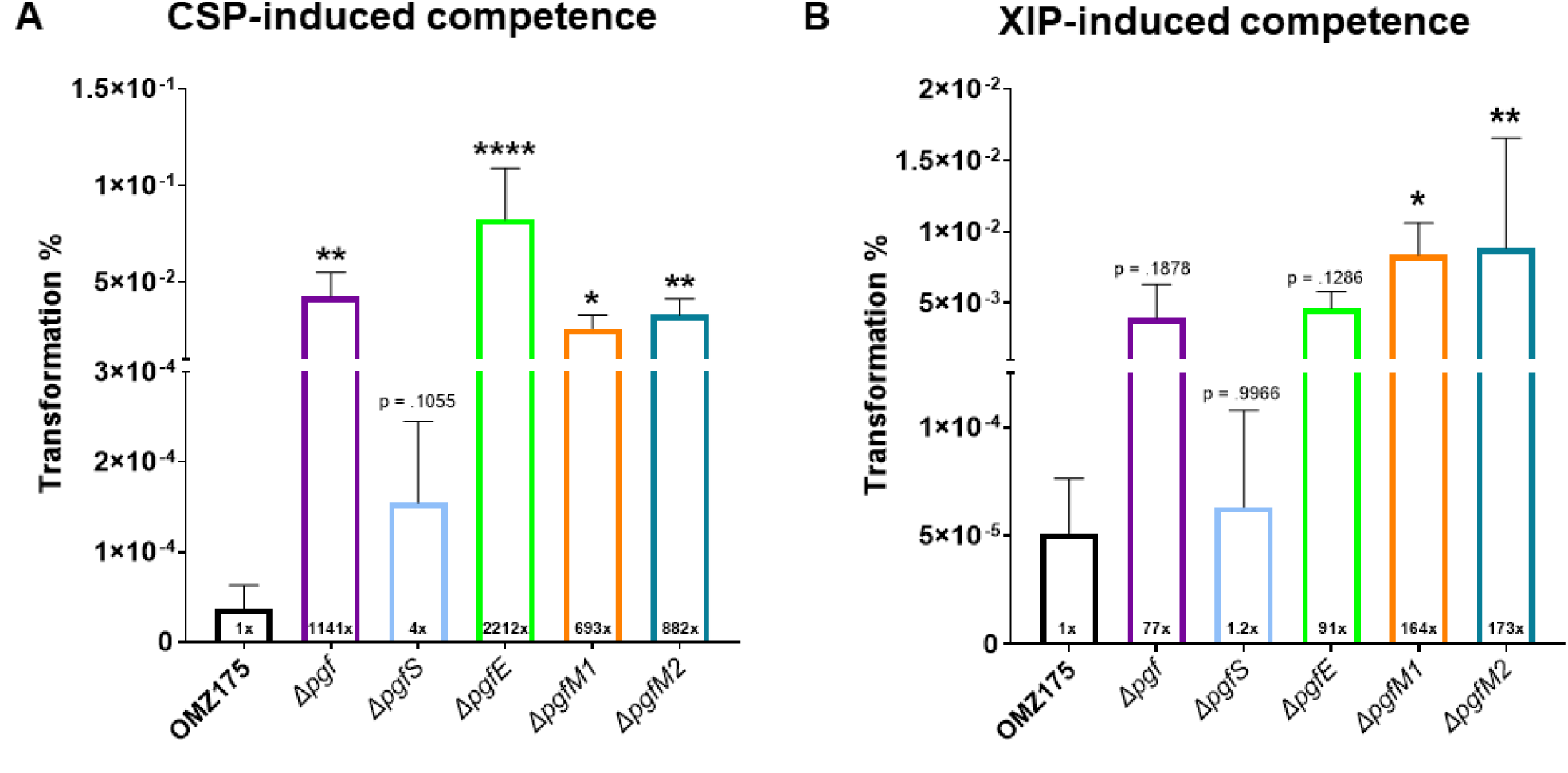
The lack of proper glycosylation status affects *S. mutans* OMZ175 competence. Transformation assays via stimulation of either *comABCDE* (A) or *comRS* (B) pathways with CSP and XIP, as indicated. The number inside each bar indicates the average fold increase in competence for each mutant in comparison to the parental strain. One-way ANOVA was utilized for comparisons, with a post-hoc test to compare differences between mutants and the parental strain. * = p < .05; ** = p < .01; **** = p < .0001. Non-significand comparisons have their p values shown above the bar. Bars indicate mean values, and error bars indicate standard deviations. N = 4.

### Deletion of the *pgf* operon significantly diminishes *S. mutans* fitness in the rat model of oral colonization

The role of the Pgf machinery in *S. mutans* OMZ175 fitness *in vivo* was analyzed using an established rat oral colonization model by infecting the animals with the parent strain or its isogenic Δ*pgf* mutant (quadruple mutant). Daily infection inocula for both strains were comparable (approximately 7.50 x 10^8^ CFU/mL), and rats from both groups had comparable weight gain throughout the entire experiment (data not shown). Cell homogenates from dissected jaws were plated on blood-agar to estimate the total oral microbiota, and on BHI supplemented with appropriate antibiotics to estimate *S. mutans* numbers. The CFU counts for blood-agar were comparable between strains, indicating that the total culturable microbiota was not affected by the infection with either type of bacteria (Figure 7A). Interestingly, a 10-fold decrease in counts recovered from the rats’ jaws infected with the *Δpgf* mutant was observed in comparison to the parent strain. When the *S. mutans* counts were represented as a percentage of the total microbiota, a significant decrease from 41.1 ± 8.7% (parental) to 16.8 ± 13.9% (*Δpgf*) in *S. mutans* percentage of total oral microbiota was observed in the *Δpgf* mutant (Figure 7B). Thus, our findings suggest that protein glycosylation by the Pgf machinery is important for oral colonization and, therefore, the overall fitness of *S. mutans* in its natural habitat. Taken altogether, the oral colonization defect of the *Δpgf* mutant is likely related to its severe biofilm defect, reduced ability to survive in saliva, and altered cell envelope homeostasis.

**Figure 7.**
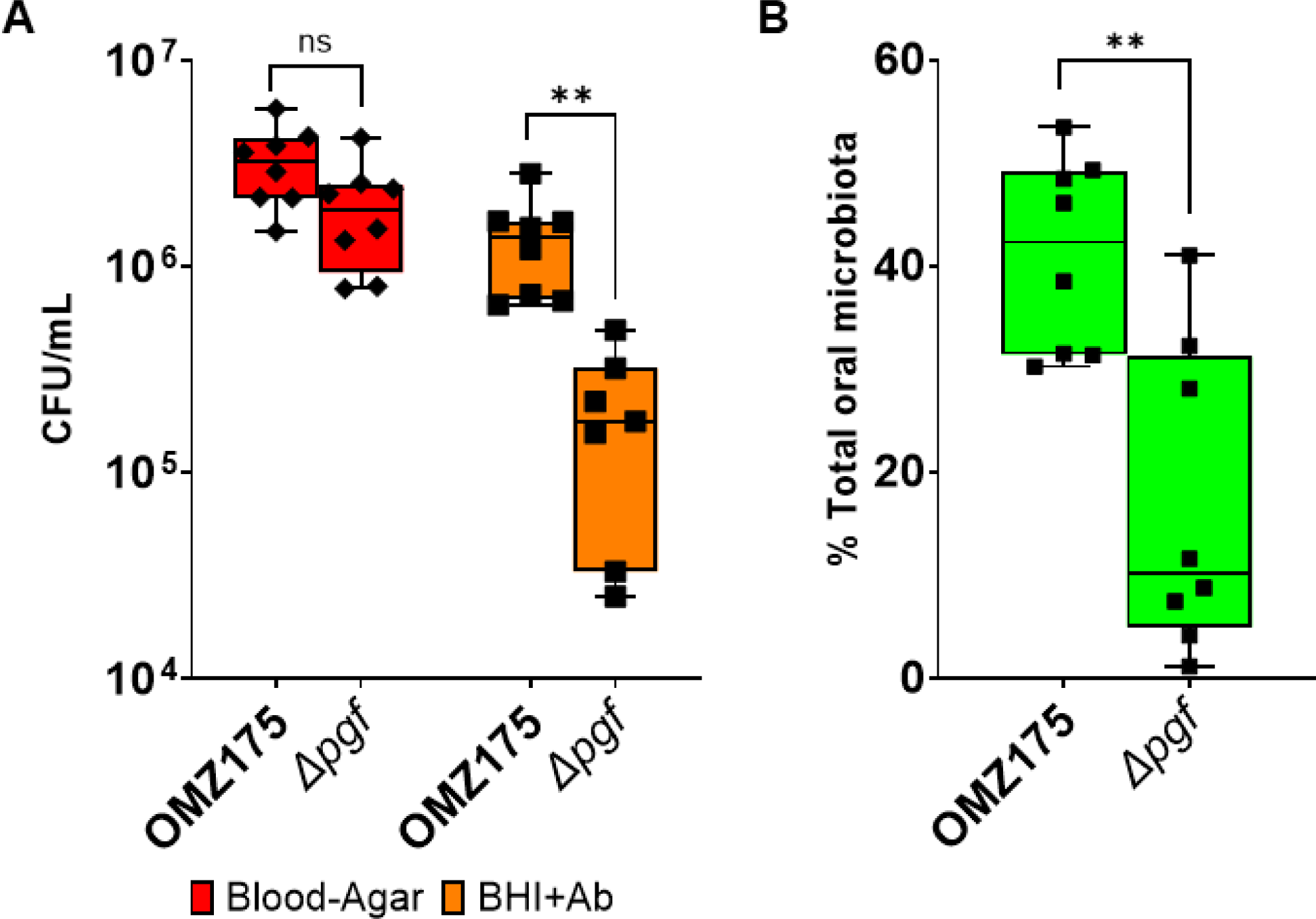
The Pgf glycosylation machinery contributes to *S. mutans* fitness *in vivo*. Rat oral colonization with *S. mutans* OMZ175 and the quadruple *Δpgf* strain. (A) Direct comparison of recovered CFUs between strains for each type of media. Blood-agar indicates CFU counts of total cultivable microbiota, and BHI+antibiotics (BHI+Ab) reveals *S. mutans* CFU counts. (B) Comparison between the percentage of *S. mutans* relative to the total oral microbiota for each strain. T-tests were performed for each pair. ns = not significant; ** = p < .01. Bars indicate mean values, and error bars indicate standard deviations. N = 8.

### Glycoproteomics analysis of the Cnm adhesin reveals HexNAc_2_ glycosylation in the parent strain and suggests competition between the Pgf machinery with the serine/threonine kinase PknB for common amino acid residues

We have recently reported that Cnm is glycosylated by the Pgf glycosylation machinery with predominantly HexNAc_2_ residues in its threonine-rich repeat region (TRRR) (Avilés-Reyes *et al*., 2018; Andresen *et al*., 2022a). Surprisingly, we discovered that the TRRR region of Cnm is phosphorylated in the absence of Pgf-mediated glycosylation (Figure 8). Figure 8A summarizes the findings from our mass spectrometry (MS) analysis. In an attempt to overcome the limitations of obtaining peptide cleavage from the highly glycosylated Cnm, we constructed a mutant harboring a truncated variant of the *cnm* gene, coding for a Cnm protein containing only the first three threonine-rich repeats (tCnm) (Andresen *et al*., 2022a). This truncated gene was inserted in the parental OMZ175 and also in the mutants *ΔpgfS* (no Cnm glycosylation) and Δ*pgfM2* (partial Cnm glycosylation). Thus, we found that the engineered tCnm obtained from the parent strain was modified with HexNAc2 in the TRRR (Figure 8B). Examination of purified native full-length Cnm from the *ΔpgfS* mutant strain showed extensive signs of phosphorylation by using the Byonic software analysis (Figure S9 and Figure S10). MS/MS fragmentation of one of those peptides confirmed extensive phosphorylation of the TRRR (Figure 8C). Similar Byonic analysis of full-length Cnm from the parent strain showed no evidence of phosphorylation (Figure S10), but the TRRR could not be appropriately detected due to extensive glycosylation. In contrast, tCnm from the *ΔpgfM2* mutant showed less glycosylation compared to the parent strain, so we tested whether Cnm from this mutant could be undergoing glycosylation and phosphorylation at the same time. Analysis of tCnm from the *ΔpgfM2* mutant (Figure 8D) demonstrated that peptides exist with one less HexNAc_2_ modification than parent strain peptides, and some do indeed possess an additional phosphate modification suggesting that both post-translational modifications can be present in this Cnm variant. We attempted to purify tCnm from the *ΔpgfS* strain for an MS analysis, but it appears to be too unstable as it quickly degrades during purification and sample processing. Future studies will examine the glycosylation and phosphorylation profiles of Cnm (and tCnm) in the other two *pgf* mutants, *ΔpgfE* and *ΔpgM1*. We acknowledge that these analyses remain challenging due to the heterogeneity observed in post-translational modifications and the instability of the truncated variants. In mammalian cells, O-linked glycosylations and O*-*linked phosphorylations were shown to crosstalk and compete for the same amino acid residues (threonines and serines), and changes in the balance between these two post-translational modifications may be implicated in an array of human pathologies like cancer and Alzheimer’s disease (Wang *et al*., 2012; Takahashi *et al*., 2022; Yi *et al*., 2022; Li *et al*., 2022). In bacteria, the crosstalk between protein phosphorylation and glycosylation remains largely underexplored. Interestingly, *S. mutans* harbors only one serine/threonine kinase, namely PknB (Banu *et al*., 2010), and future studies will include mutants for this gene in parental and *pgf* mutant backgrounds to further characterize the post-translational modification profile of the model protein Cnm. A better understanding of the Pgf substrates and specificities will be crucial for a deeper understanding of the phenotypes observed in our panel of mutant strains. For instance, the presence of heavily phosphorylated Cnm in the cell surface of *ΔpgfS* may contribute to its significantly more negative surface charge when compared to the parent strain (Figure 5A).

**Figure 8.**
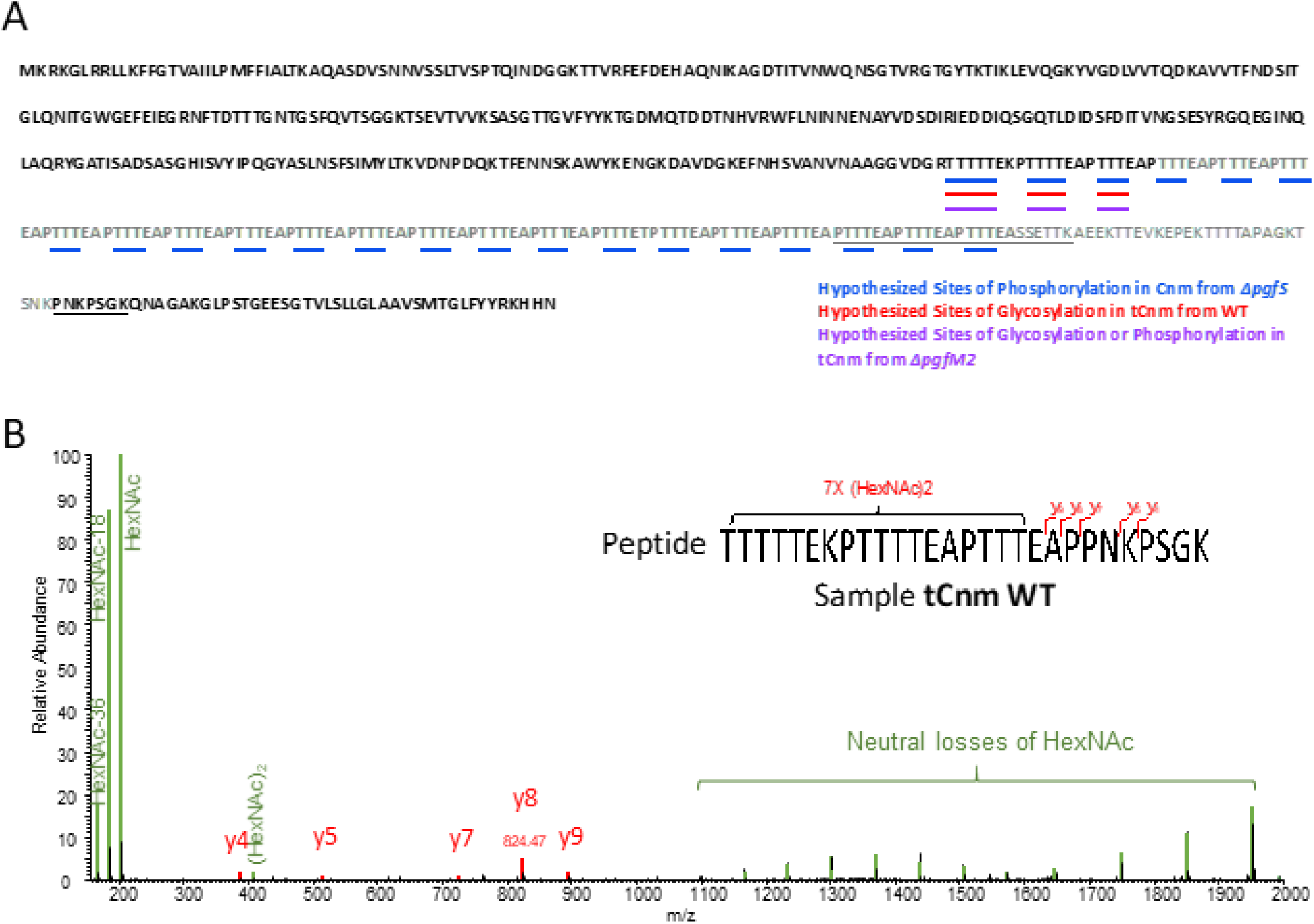
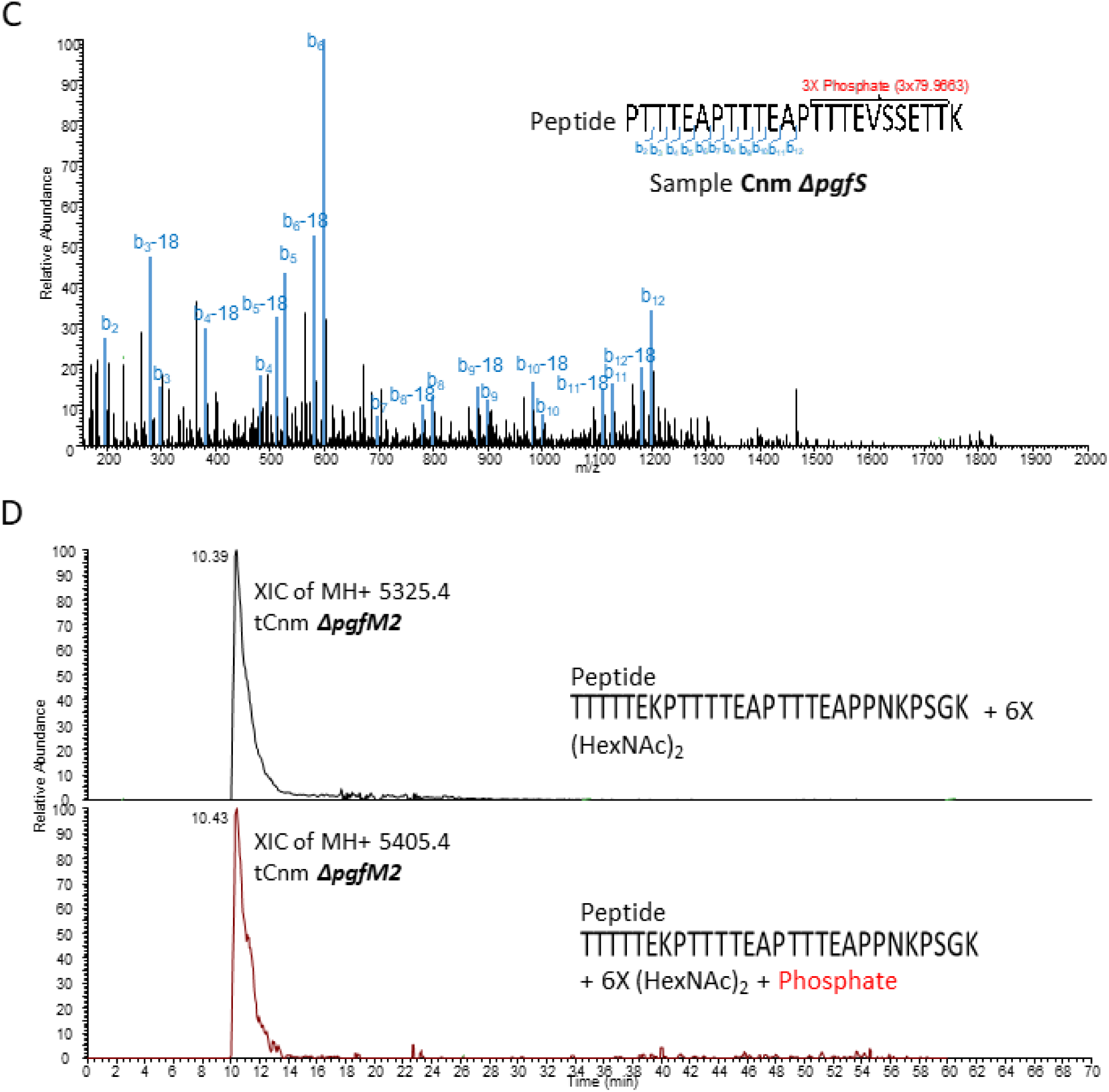
Cnm and the truncated Cnm (tCnm) are glycosylated with HexNAc in the threonine-rich repeats whereas the unglycosylated Cnm and tCnm from the *ΔpgfS* mutant are phosphorylated. Mass spectrometric analysis of glycosylation and phosphorylation status of Cnm and tCnm purified from parent and *ΔpgfS and ΔpgfM2* mutants. (A) Sequence of Cnm (full sequence shown in black and grey) and tCnm (sequence shown in black only) with proposed sites of post-translational modifications within the threonine-rich repeat region. The peptides shown in panels B-D are indicated as underlined. (B) MS/MS fragmentation (HCD) of a phosphorylated peptide from Cnm purified from the *ΔpgfS* mutant. (C) MS/MS fragmentation (HCD) of a glycosylated peptide from parent tCnm. (D) Extracted ion chromatographs (XICs) of peptides from tCnm purified from *ΔpgfM2* indicating partial glycosylation and partial phosphorylation.

## Conclusions

The Pgf machinery regulates, through post-translational protein glycosylation and possibly by modifying other targets like the RGP, a wide range of complex biological processes in *S. mutans* that are essential for its pathobiology (Table 1, with a visual representation in Figure 9) such as proper biofilm development, survival in saliva, cell envelope homeostasis, competence and, most importantly, oral colonization *in vivo*. Our findings suggest functional redundancies between the Pgf machinery, especially PgfS, and the RGP synthesis pathways. We also found that in the absence of proper glycosylation through the Pgf machinery, Cnm becomes phosphorylated in the TRRR, suggesting that Pgf machinery and the only serine-threonine kinase present in *S. mutans* compete for the same sites in Cnm.

**Figure 9.**
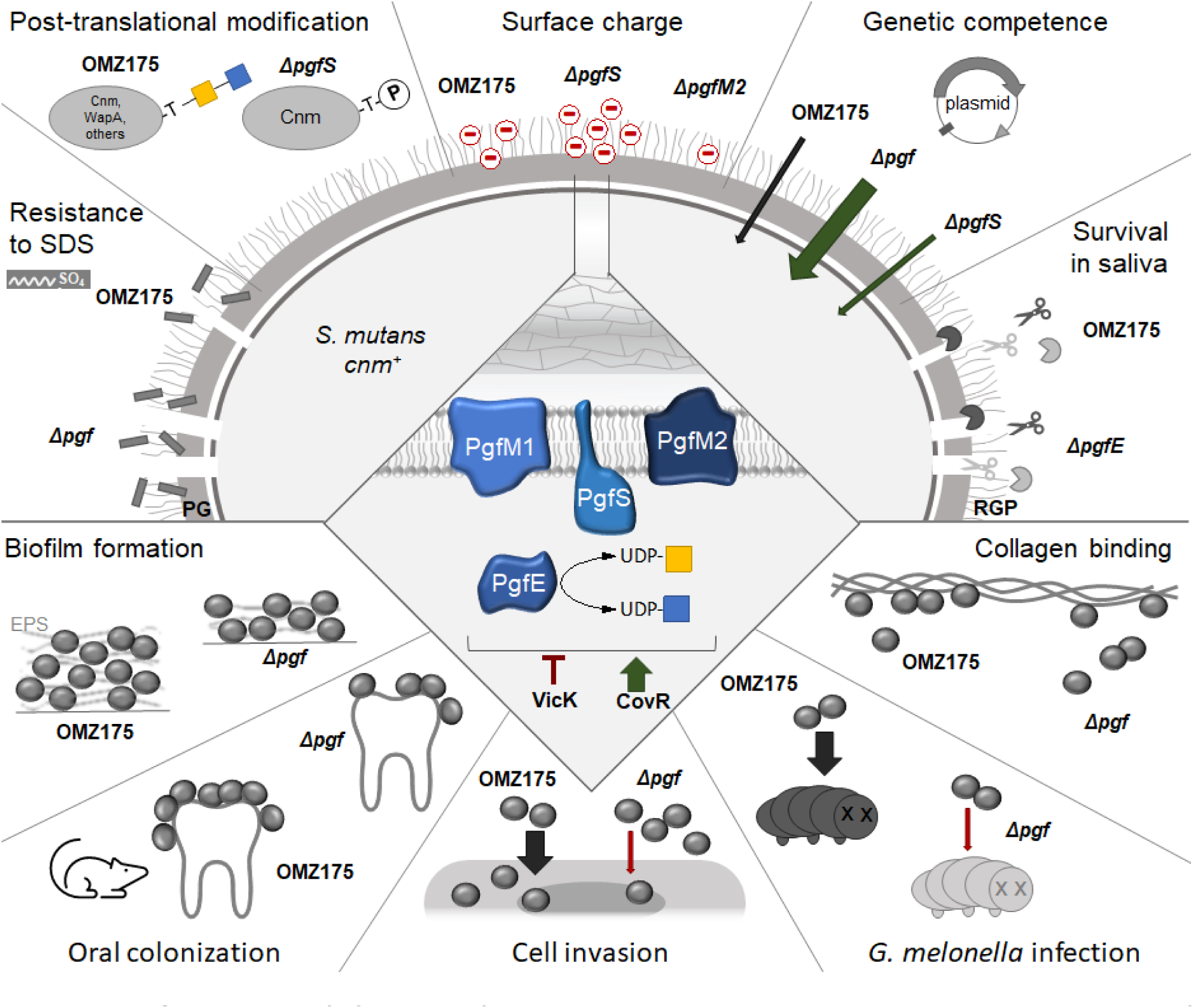
Summary of findings from our current and previous studies. The Pgf glycosylation machinery is directly involved in the glycosylation of surface adhesins and possibly of the rhamnose-glucose polysaccharide layer. Glycosylation appears to be favored over phosphorylation, suggesting crosstalk or competition between post-translational modification pathways. Glycosylation also modulates surface charge, membrane homeostasis, genetic competence, biofilm formation, saliva survival, and fitness in an oral colonization model. Previously studied phenotypes associated with proper glycosylation of Cnm are endothelial and epithelial cell invasion, systemic infection in the *Galleria mellonella* model, and collagen binding. Expression of *pgf* genes is under positive regulation via CovR and negative regulation via VicRKS. PG = peptidoglycan; RGP = rhamnose-glycose polysaccharide; P = phosphate; EPS = extracellular polysaccharides. Following the Symbol Nomenclature For Glycans, GlcNAc (N-Acetylglucosamine) is depicted as a blue square and GalNAc (N-Acetylgalactosamine) is depicted as a yellow square.

## Material and Methods

### Bacterial strains and growth conditions

The strains used in this study are listed in Table 2. All *pgf* mutants have been previously complemented (Avilés-Reyes *et al*., 2014a; Avilés-Reyes *et al*., 2018). *S. mutans* was routinely grown in Brain Heart Infusion (BHI) media and Chemically Defined Media (CDM) (van de Rijn and Kessler, 1980) with 1% (m/v) glucose at 37°C and 5% CO_2_. When required, Kanamycin or Erythromycin was added to the media at 1 mg.mL^-1^ or 0.3 mg.mL^-1^, respectively. Biofilms were grown in CDM with 1% (m/v) sucrose as a carbon source.

**Table 2.** Bacterial strains utilized in this study.

| Strain | Genotype | Reference |
| --- | --- | --- |
| *Streptococcus mutans*<br>OMZ175 | Parental strain | B. Guggenheim |
| | $\Delta gtfA$ *Erm*<sup>R</sup> | (Miller *et al.*, 2015) |
| | $\Delta pgfS$ *Kan*<sup>R</sup> | (Avilés-Reyes *et al.*, 2014a) |
| | $\Delta pgf$ *Kan*<sup>R</sup> | (Avilés-Reyes *et al.*, 2018) |
| | $\Delta pgfE$ *Kan*<sup>R</sup> | |
| | $\Delta pgfM1$ *Kan*<sup>R</sup> | |
| | $\Delta pgfM2$ *Kan*<sup>R</sup> | |
| | $\Delta vicK$ *Kan*<sup>R</sup> | (Alves *et al.*, 2018) |
|  | *tcnm* (no resistances) | (Andresen *et al.*, 2022a) |
| | *tcnm* $\Delta pgfS$ *Kan*<sup>R</sup> | This study |
| | *tcnm* $\Delta pgfM2$ *Kan*<sup>R</sup> | This study |

### Bacterial mutagenesis

An OMZ175 mutant harboring a truncated *cnm* gene that generates a protein containing only the first three threonine-rich repeats in its B domain was generated as described elsewhere (Andresen *et al*., 2022a). This mutant strain (*tcnm*) had either the *pgfS* or the *pgfM2* genes deleted by the same strategy used to create the *ΔpgfS* and *ΔpgfM2* mutants (Avilés-Reyes *et al*., 2018), generating the double mutants *tcnmΔpgfS* and *tcnmΔpgfM2*.

### Biofilm biomass assays

Quantification of biofilm biomass and colony-forming unit counts (CFUs) were performed as previously described with some modifications (Lemos *et al*., 2010). Briefly, overnight cultures of *S. mutans* strains grown in BHI were diluted 1:100 in CDM containing 1% sucrose as the sole carbon source in 96-well plates. Four biological replicates were grown in technical triplicates on duplicate plates for 48 hours. Media was carefully removed, and wells were washed twice with sterile PBS to remove non-adherent cells. Biofilms were scraped from one of the plates, resuspended in 20 µL of PBS, and plated on BHI-agar plates for determination of CFUs. Biofilms of the second plate were stained for 15 minutes with a 0.05% crystal violet solution. Wells were washed twice with PBS and stained biofilms were resuspended in 7% acetic acid (v/v) and absorbance was read at 595 nm in a Synergy H1 hybrid multimode reader (BioTek).

### Confocal Laser Scanning Microscopy

Overnight cultures were diluted 1:100 in CDM + 1% sucrose in µ-Slide 8 Well polymer coverslip chambers (Ibidi) and grown for 48 hours. Spent media and planktonic cells were carefully removed by pipetting and biofilms were gently washed twice with a sterile 0.85% (v/v) NaCl solution. Then, a 0.85% NaCl solution containing 5 µM Syto9 (ThermoScientific) to identify live *S. mutans* cells and 1 µM Dextran Alexa Fluor 647 (ThermoScientific) to identify extracellular polysaccharides was added to stain biofilms for 30 minutes. Biofilms were again washed to remove unbound dyes, and images were acquired using a Nikon Ti2 Confocal Microscope with a Plan Apo λ 60x oil objective coupled with a Nikon C2 Plus camera. NIS-Elements software (Nikon) was used to analyze three-dimensional images with a 1 µm height difference between z-stacks for three spots of each biofilm using at least 4 biological replicates. Thresholds were determined for the creation of binary layers for *S. mutans* cells and extracellular polysaccharides, allowing for quantitative analysis of biofilms (Paula *et al*., 2020).

### Cnm purification and digestion

Cnm from OMZ175 and the *ΔpgfS* mutant, and tCnm from OMZ175 parental background and the *tcnmΔpgfS* and *tcnmΔpgfM2* mutant backgrounds were purified by affinity chromatography using custom columns coupled with anti-Cnm collagen-binding domain (CBD) antibodies, as previously published (Avilés-Reyes *et al*., 2014a; Avilés-Reyes *et al*., 2014b). Briefly, the OMZ175*ΔvicK* strain, known to overexpresses Cnm (Alves *et al*., 2018), and the *ΔpgfS* mutant were grown overnight in liquid cultures and then lysed by bead beating in the presence of Halt protease inhibitor single-use cocktail (Thermo Fischer Scientific) and ethylenediaminetetraacetic acid (EDTA). The soluble protein fractions were then bound to custom N-hydroxysuccinimide (NHS)-anti-CBD columns with overnight rocking at 4°C. Different columns were used for proteins from different strains to avoid cross-contamination. Then, columns were washed with 15 mL of 1x PBS at pH 7.2, and protein was eluted by incubation in a 0.1 M glycine buffer (pH 2.5) for 5 min. Elutions were immediately neutralized with 1/10 volume of a basic 1 M Tris buffer (pH 8.0). Purified native proteins were later concentrated and dialyzed against a 50 mM ammonium bicarbonate solution. Proteins were then digested in solution as described elsewhere (Shajahan *et al*., 2017). Although there are no cysteines in Cnm to form disulfide bonds, dithiothreitol (DTT) was used at a low concentration (5mM) to help disrupt the protein tertiary structure. To this, sequencing grade trypsin (Promega) was added at a protease:protein mass ratio of 1:20 and digested overnight at 37°C. The enzymatic digestion was halted the next day by briefly heating at 100°C, and the digested peptides were cleaned-up and dried by C18 solid phase extraction cartridges (Hawach Scientific) and dried on a vacuum centrifuge.

### Mass spectrometry analysis

Tandem liquid chromatography mass spectrometry (LC-MS/MS) was performed on an Orbitrap Eclipse Tribrid mass spectrometer equipped with a nanospray ion source coupled with a Thermo Ultimate RSLCnano chromatography system (ThermoFisher). Digested peptides were reconstituted in 0.1% (v/v) formic acid (FA) and pre-filtered through a 0.2 µm spin filter before injection into the system. A commercial nano-LC column (ThermoFisher) of 15 cm length with 75 μm internal diameter and filled with 3 μm C18 material (reverse phase) was used for the chromatographic separation of samples. The precursor ion scans were acquired at 120,000 resolution in the Orbitrap analyzer, and precursors at a time frame of 3 seconds were selected for subsequent MS/MS higher-energy C-trap dissociation (HCD) fragmentation in the Orbitrap analyzer at 15,000 resolution. Runs of each digest were conducted for 70 or 180 min for phosphorylation and glycosylation analysis, respectively. Solutions of 0.1% FA and 80% acetonitrile–0.1% FA were used as mobile phases to separate the glycopeptides. The threshold for triggering an MS/MS event was set to 1,000 counts, and monoisotopic precursor selection was enabled. Data were processed with Byonic (Protein Metrics v4.0.12) software using the full-length sequence of Cnm as a reference. Precursor mass tolerance was set to 5 ppm and fragment mass tolerance was set to 20 ppm. Modifications included oxidation and deamidation on methionine and asparagine respectively, and phosphorylation on serine, threonine, and tyrosine. Detection of multiple phosphates (e.g., 2X phosphorylation = 159.9327) was also programmed for analysis.

### Zeta potential measurement

Estimation of *S. mutans* envelope charge via zeta potential in liquid culture was performed as previously described with some modifications (Wilson *et al*., 2001; Soni *et al*., 2008; Ng and Ting, 2016). Cultures were grown in BHI to an OD_600_ of 0.15, then cells were harvested by centrifugation (10,000 x g for 2 minutes) and washed three times with 0.1X PBS to remove loosely bound ions at the bacterial surface. Pellets were resuspended in 0.1X PBS to prevent osmotic stress and cell clumping. Then, 800 µL of each cell suspension was added to a 12 mm glass cuvette (Malvern Panalytical) which was closed with a Dip Cell with palladium electrodes with 2 mm spacing (Malvern Panalytical), inserted into a Malvern Zetasizer Ultra particle analysis system (Malvern Panalytical), and equilibrated to 25°C for 60 seconds. Readings were performed using 30 individual measurements with self-refinements, resulting in sharp and stable peaks for each sample.

### Minimum Inhibitory Concentration determination

To determine the MIC of SDS and tunicamycin, overnight cultures were used to respectively inoculate BHI supplemented with up to 0-0.01% (m/v) Sodium Dodecyl Sulphide or up to 20 µg/mL of tunicamycin from *Streptomyces lysosuperficus* (Millipore Corp) solubilized in Dimethyl Sulfoxide, in quadruplicate. After 24h of incubation at 37°C in 5% CO_2_, the OD_600_ was read using a Synergy H1 hybrid multimode reader (BioTek). MIC values were considered as the concentration in which >90% of inhibition occurred.

### Ethidium Bromide permeability assays

The assay was performed as previously described, with minor modifications (Patry *et al*., 2019). Briefly, *S. mutans* strains were grown to mid-logarithmic phase (OD_600_ of ≈0.5) in BHI, harvested by centrifugation, and resuspended in PBS to an OD_600_ of 0.2. The bacterial suspensions were mixed with 20 µM ethidium bromide (EtBr) solution to a final concentration of 1 µM. Immediately after the addition of the EtBr, spectrophotometric monitoring at 530 nm extinction and 600 nm emission was started using a Synergy H1 hybrid multimode reader (BioTek) with measurements at 20-second intervals. At least four independent biological replicates were performed for each strain, with two technical replicates.

### Antibiotic resistance disc assays

Antibiotic discs were prepared using sterile Whatman paper discs (7 mm in diameter) saturated with 20 µL of antibiotic stock solutions (Penicillin 1.5 mg.mL^-1^, Vancomycin 1.5 mg.mL^-1^, Gentamycin 1.5 mg.mL^-1^, Streptomycin 1.5 mg.mL^-1^, Chloramphenicol 1.5 mg.mL^-1^, Tetracycline 1.5 mg.mL^-1^, Ampicillin 0.5 mg.mL^-1^). The discs were placed on BHI agar plates which had been freshly coated with 150 µL of mid-logarithmic phase *S. mutans* cultures. Diameters of zones of growth inhibition around each antibiotic disc were measured after 24 hours of incubation. Assays were performed with three biological replicates per strain.

### Opsonophagocytosis assays

The method for the opsonophagocytosis assay was adapted from a previously published method (Andresen *et al*., 2022b). Blood was drawn from healthy, adult, human volunteers at the Clinical Translational Research Unit of the University of Georgia by venipuncture under informed consent. Heparin was used as an anticoagulant. Briefly, 5 mL of whole human blood was layered onto 5 mL of Histopaque-1077 Hybri-MaxTM (Sigma Life Science, H8889-100 mL) in a column, followed by centrifugation at 350 x g for 30 min to induce the formation of a density gradient. The peripheral blood mononuclear cells (PBMCs) were collected from the corresponding layer, washed in PBS, and resuspended in Dulbecco’s Modified Eagle Medium (DMEM). PBMCs were stained with Methylene Blue and counted by visualization of intact cells in a hemocytometer. Briefly, *S. mutans* cultures were grown to mid-exponential phase (OD_600_ of ≈0.5) and adjusted to an OD_600_ of 0.01 in DMEM (8×10^6^ CFU/mL). Then, 1.25×10^5^ CFU of bacteria were combined with 5×10^5^ PBMCs for a final multiplicity of infection (MOI) of about 1:4 bacteria to PBMCs. Pooled normal human complement serum (NHS; a source of complement) (Innovative Research) or heat-inactivated NHS (HIS) were added to the bacterial:PBMC mixture in DMEM to a final volume of 200 µL. As a control condition, the complement components were heat inactivated by heating to 56 °C for 1 h. The components were incubated for 2 hours, then serially diluted and plated on BHI for CFU quantifications of viable bacterial cells. The percentage of survival was calculated relative to controls free of PBMCs (= 100% survival).

### Serum survival assay

Pooled human complement serum was purchased from Innovative Research. The serum survival assay was done as previously described (Lees-Miller *et al*., 2013) with some modifications. *S. mutans* strains grown to mid-exponential phase were harvested by centrifugation and adjusted to an OD_600_ of 2. In a 96-well plate, 5 µL of bacterial suspension were combined with 20 µl (20% v/v final concentration) of baby rabbit complement (BRC) and with heat-inactivated (1 h at 56 °C) normal human serum (Innovative Research) in a final volume of 100 µL. The mixtures were incubated for 24 or 48 hours before plating serial dilutions onto BHI plates. The survival of *S. mutans* in the presence of BRC was calculated as a percentage of survival in heat-inactivated BRC (1 h at 56 °C). Four biological replicates of the experiment were performed for each strain. Controls for the assay were performed using the *Acinetobacter baumannii* 5075 WT (robust survival) and *A. baumannii* 5075*ΔpglC* (complete killing) (Crippen *et al*., 2021).

### Saliva survival assay

Saliva was pooled from healthy donors and clarified by filtration (0.2 µm cutoff filter). Stationary phase *S. mutans* cultures were harvested by centrifugation and resuspended in PBS to one-tenth of the original volume. Twenty microliters of each concentrated culture were added to 200 µL of pooled human saliva supplemented with 20 µM of glucose and incubated for 48 hours. Samples were removed at various time points to allow for the counting of viable CFUs.

### Genetic competence

Biological replicates (three for each strain) of *S. mutans* were subcultured into BHI or CDM + 1% glucose and incubated until the cultures reached an OD_600_ of approximately 0.08. The BHI-grown cultures were then supplemented with 1 µM Competence Stimulating Peptide (CSP; SGSLSTFFRLFNRSFTQALGK) and the CDM-growth cultures were supplemented with *comX*-Inducing Peptide (XIP; GLDWWSL) for the stimulation of early competence via the *comABCDE* and *comRS* pathways, respectively. Five hundred nanograms of purified spectinomycin-resistance conferring plasmid pDL278 (Addgene, #46882) were also added to each culture. Incubation was continued for four hours, followed by tenfold serial dilutions plated on BHI-agar plates with or without spectinomycin (1 mg/mL). After 48 hours, CFUs were enumerated, and transformation efficiency was calculated by dividing the CFUs counted from the plates with antibiotic by the CFUs counted from the plates without antibiotic.

### Rat oral colonization assay

An established rat oral colonization model (Miller *et al*., 2015; Galvão *et al*., 2017) was performed with minor modifications using the parental OMZ175 strain and a *Δpgf* mutant. An *S. mutans* OMZ175 strain harboring an Erythromycin resistance cassette integrated at the *gtfA* locus was used as the parent strain in this study (Miller *et al*., 2015). This gene has been demonstrated not to interfere with *S. mutans* virulence in rat oral colonization models (Barletta *et al*., 1988), and therefore this mutant is herein referred to as the parental strain. Upon arrival, Sprague-Dawley female rats were provided with sulfamethoxazole (0.8 mg/mL) and trimethoprim (0.16 mg/mL) in their drinking water to suppress their endogenous microbiota, facilitating infection. Antibiotics were removed three days before the animals were infected for four consecutive days by inoculating the oral cavities with cotton swabs saturated with overnight cultures of *S. mutans*. At the start of the infection period, animals were given a balanced rodent diet containing 12% sucrose (Envigo), provided for the duration of the experiment. Fourteen days after the initial oral infection, animals were sacrificed, and the lower jaws were removed for sonication in sterile PBS to remove biofilms from the tooth surfaces and to homogenize the cells. The sonicate was then tenfold serially diluted and plated on blood-agar (total bacteria), and BHI containing either Kanamycin (*Δpgf* only) or Erythromycin (Parental only). The number of total bacteria and *S. mutans* recovered is expressed as CFU per mL of sonicate on each media, and the percentage of *S. mutans* was estimated based on the number of bacteria in BHI + Antibiotic plates relative to blood-agar CFUs.

### Bioinformatics

Protein structure predictions were performed using AlphaFold2 (Jumper *et al*., 2021) in the University of Florida’s supercomputer HiPerGator 3.0. Domain function predictions were performed using InterPro (Blum *et al*., 2021), and the position of domains in transmembrane proteins was predicted using TMHMM 2.0 (Krogh *et al*., 2001). Homomer predictions were performed by template-based modeling using GalaxyHomomer (Baek *et al*., 2017) and the predicted AlphaFold2 structures of PgfS and PgfE. Protein alignments were performed using the Basic Local Alignment Search Tool (BLAST), available at the National Center for Biotechnology Information (NCBI) website (https://blast.ncbi.nlm.nih.gov/Blast.cgi).

### Statistical Analyses

Statistical analyses were performed using GraphPad Prism 9.4 (GraphPad Software, La Jolla, CA, USA). Quantitative analysis of mutant strains’ phenotypes was performed using an analysis of variance (ANOVA) with a Dunnett post hoc test, using the parental strain as a control. Comparisons between strain pairs were performed using a Student’s t-test when appropriate. A p-value of < .05 was considered to indicate a significant difference in all tests.

## Acknowledgments

We thank Dr. Balazs Rada for providing the human blood samples, and also thank the healthy volunteers for the donation of their blood and the staff of the UGA Clinical and Translational Research Unit for drawing blood used in studies, supported by the National Institutes of Health (5R01HL136707 and R21AI147097 to Balazs Rada). The authors also acknowledge the University of Florida Research Computing for providing computational resources and support that have contributed to the research results reported in this publication. URL: http://researchcomputing.ufl.edu.

## Author contributions

Conceptualization, Project administration, Resources, and Supervision: JA, JAL, CS, PA. Data Curation: NMC, SA, SAH. Formal analysis: NMC, SA, SAH, AMP. Funding acquisition: JA, JAL. Investigation: NMC, SA, SAH, AMP, TG, JKK, BAG, SS, IS. Methodology: NMC, SA, SAH, TG, JKK, BAG, IS, SS, JA, JAL, PA, CS. Validation: NMC, SA, SAH, IS. Visualization: NMC, SA, CS, JA, JAL. Writing – original draft: NMC, SA, JA, CS, JAL. Writing – review & editing: NMC, SA, SAH, BAG, AMP, TG, SS, IS, JKK, PA, CS, JA, JAL.

## Data availability statement

All data that support the findings of this study are available in the main text and the supplemental material.

## Ethics statement

Rat oral colonization protocol has been approved by the University of Florida Institutional Animal Care and Use Committee (Protocol 201810421). Human saliva was collected from healthy subjects with written consent, with approval from the University of Florida’s Internal Review Board (Protocol 01600877). Whole fresh human blood was drawn from adult volunteers at the Health Center or the Clinical Translational Research Unit of the University of Georgia under informed consent according to procedures approved by the Institutional Review Boards at the University of Georgia (Protocol 20121076906).

---

**Figure S1.**
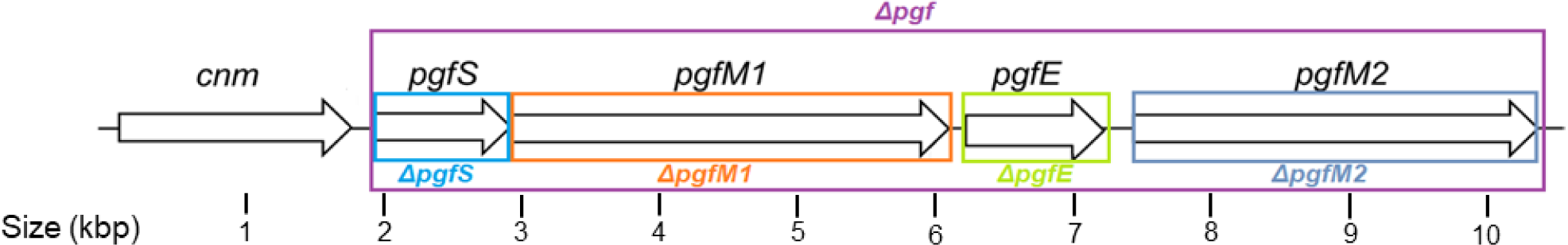
Organization of the *cnm-pgf* locus in *S. mutans* strain OMZ175, with each mutant in the panel of strains indicated in colored rectangles.

**Figure S2.**
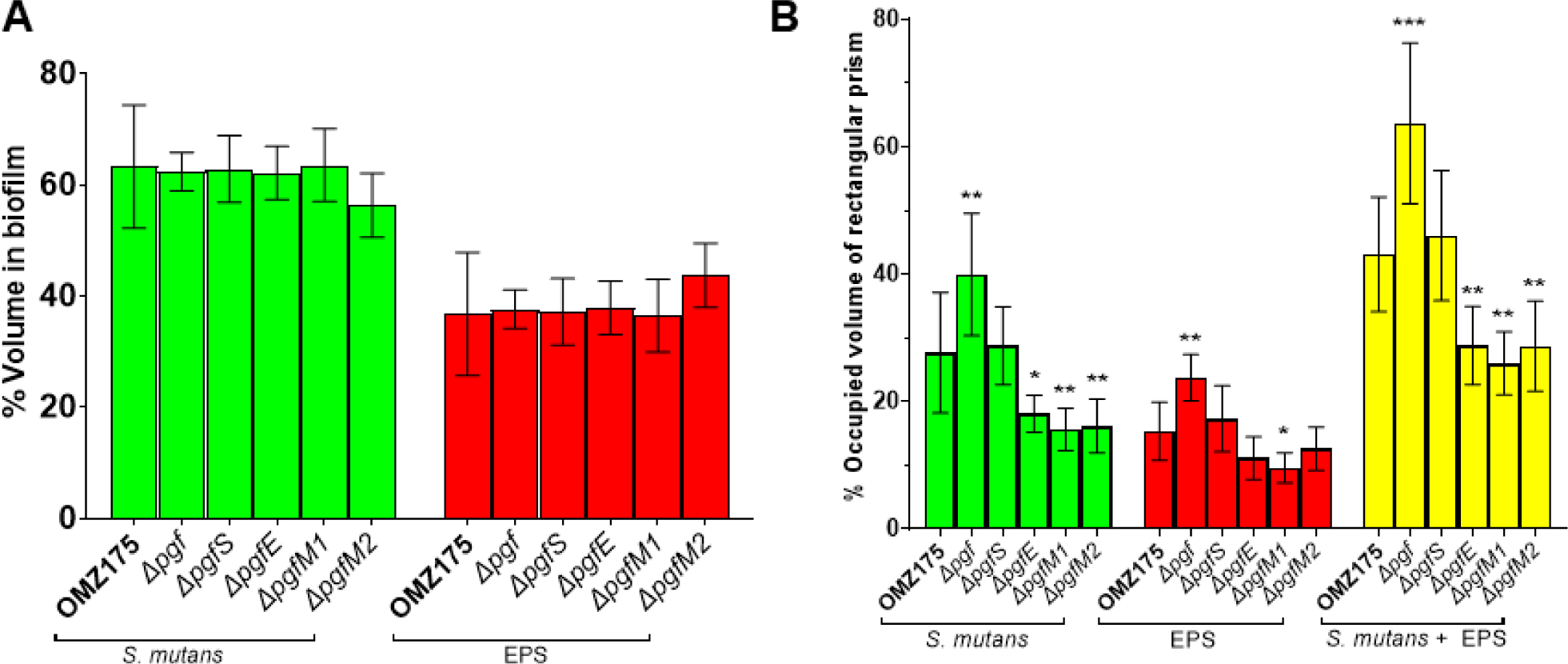
Quantitative analysis of Confocal Lases Microscopy Scanning images. (A) Distribution of *S. mutans* cells and EPS as a percentage of the volume of the sum (p = .7506). (B) Biofilm compactness as estimated by calculating the volumetric occupancy of each biofilm component relative to the rectangular prism that encases the entire biofilm (base area = 210 x 210 µm^2^, height = distance between bottom and top of biofilm) (p < .0001 for *S. mutans*; p = .0002 for EPS; p < .0001 for sum). Bars indicate means, and error bars indicate standard deviation.

**Figure S3.**
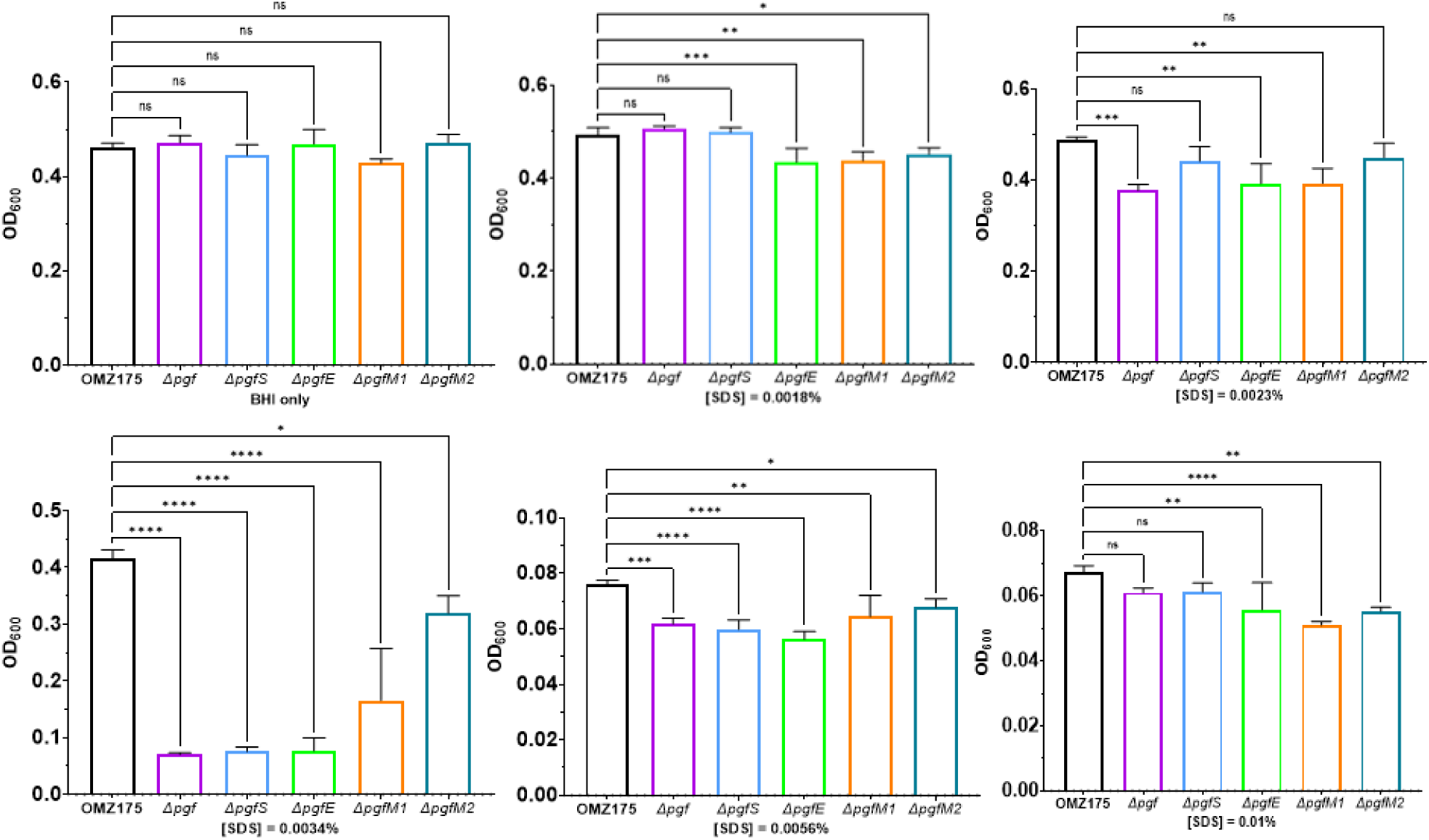
One-way ANOVA comparisons between the growth of the parental strain and each mutant in different concentrations of SDS. N = 4. ns = non-significant; * = p < .05; ** = p < .01; *** = p < .001; **** = p < .0001.

**Figure S4.**
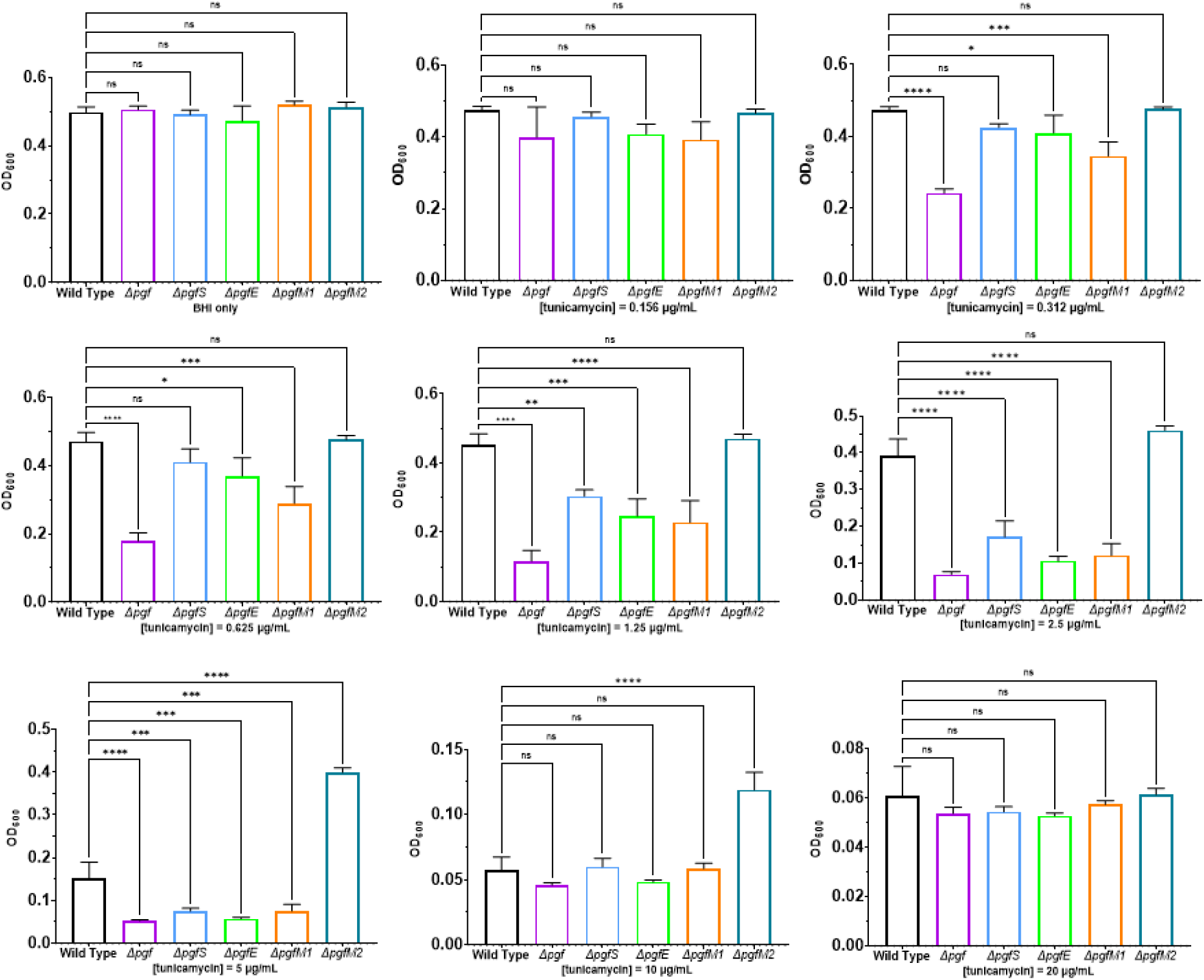
One-way ANOVA comparisons between the growth of the parental strain and each mutant in different concentrations of tunicamycin. N = 4. ns = non-significant; * = p < .05; ** = p < .01; *** = p < .001; **** = p < .0001.

**Figure S5.**
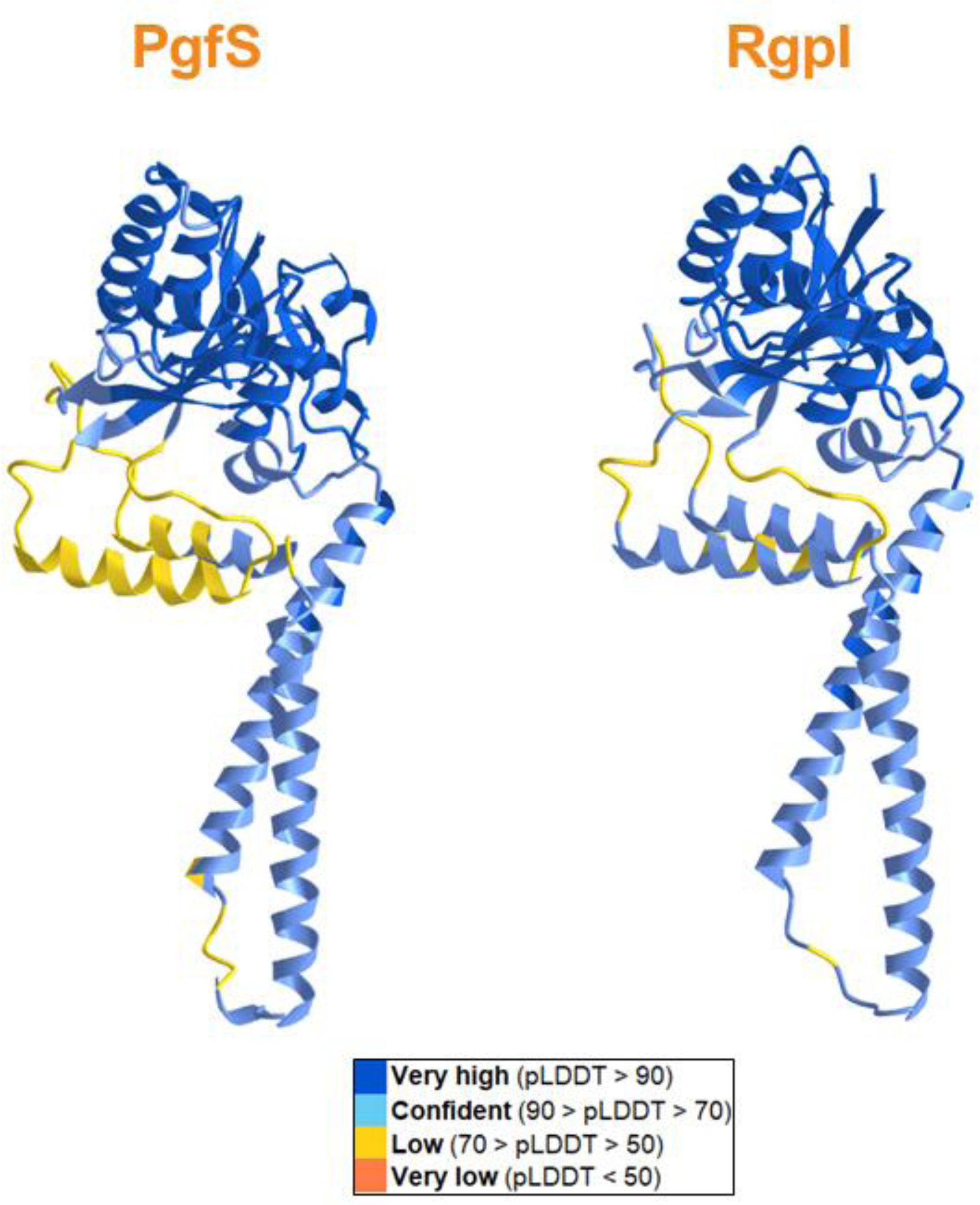
AlphaFold2 structure predictions for PgfS and RgpI from *S. mutans* indicate a high level of structure homology between the two proteins.

**Figure S6.**
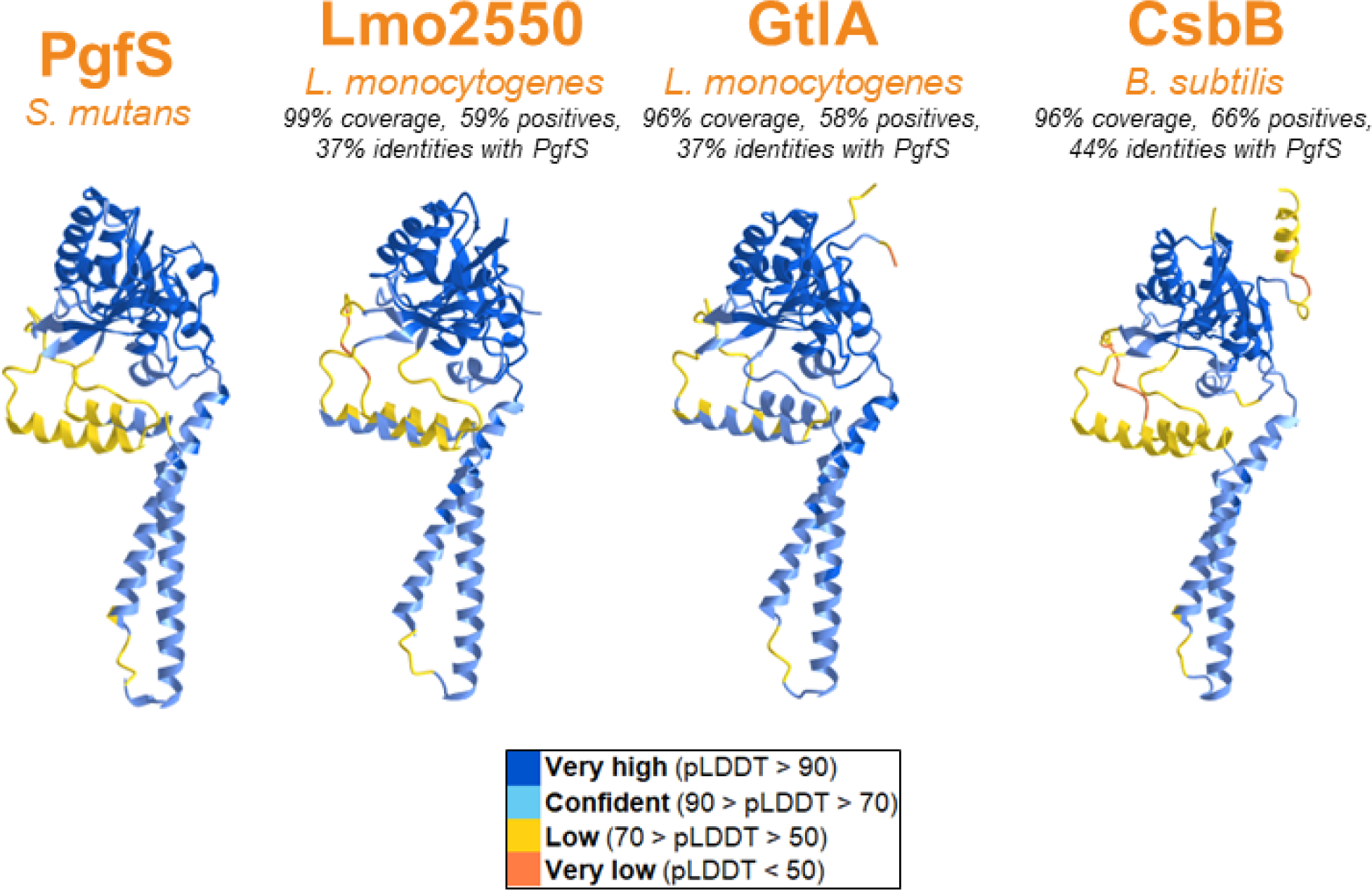
AlphaFold2 structure predictions for PgfS from *S. mutans* and PgfS-like enzymes involved in lipid glycosylation in *L. monocytogenes* and *B. subtilis*.

**Figure S7.**
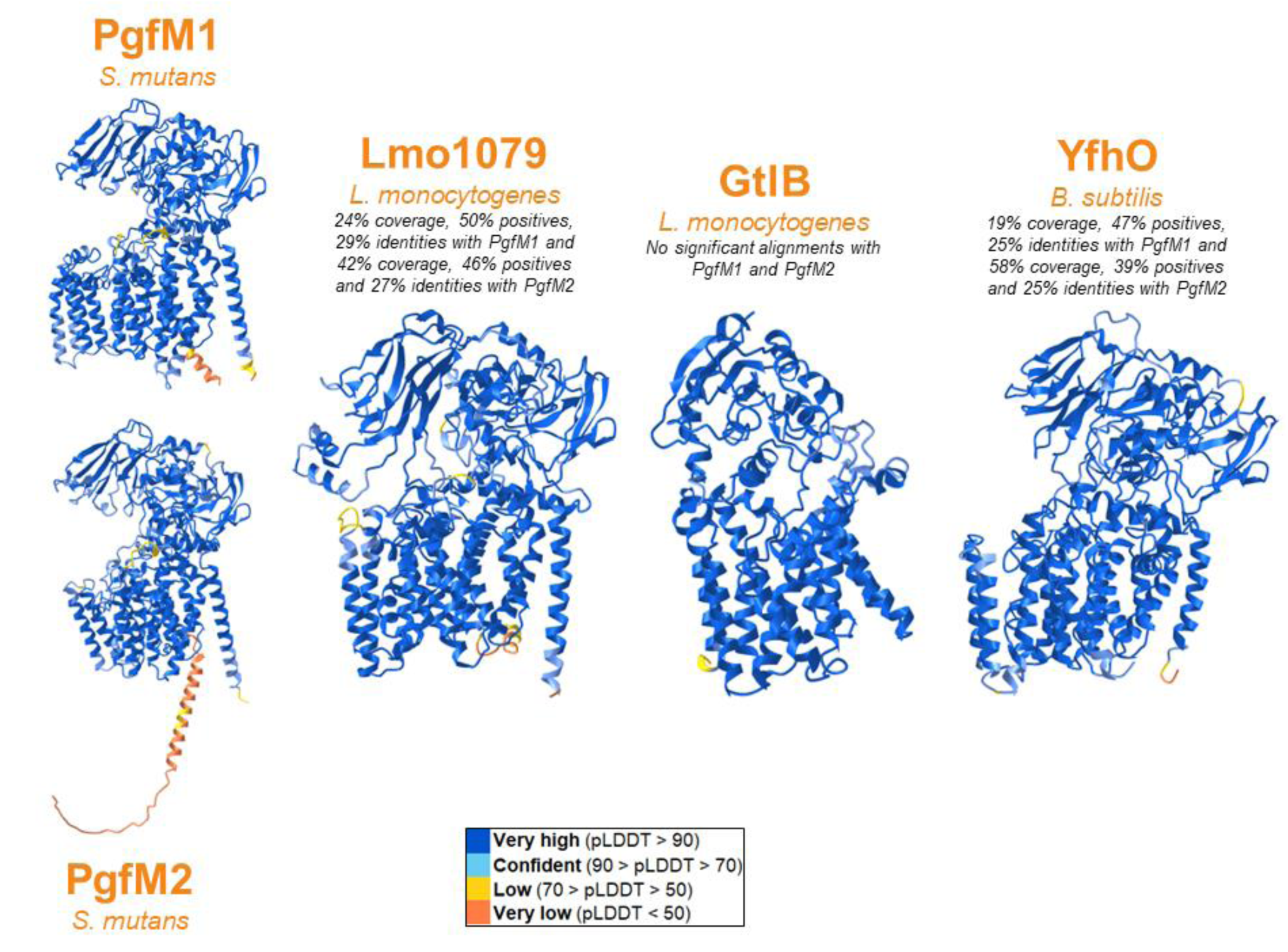
AlphaFold2 structure predictions for PgfM1 and PgfM2 from *S. mutans* and PgfM1/M2-like enzymes involved in lipid glycosylation in *L. monocytogenes* and *B. subtilis*.

**Figure S8.**
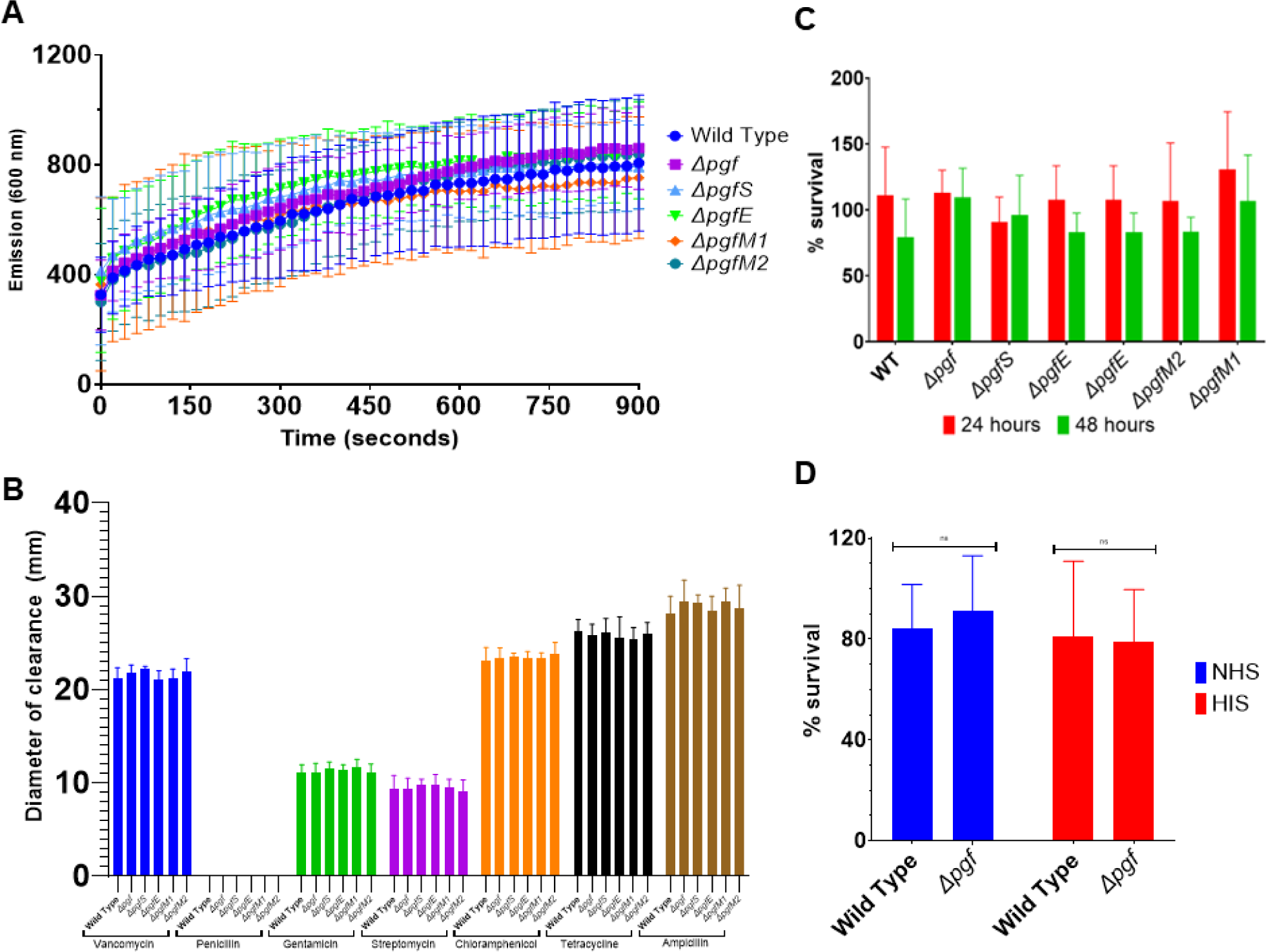
Ethidium bromide permeability (A), serum survival (B), antibiotic disc resistance (C) and opsonophagocytosis (D) assays did not reveal different phenotypes between the parental strain and the *pgf* mutants.

**Figure S9.**
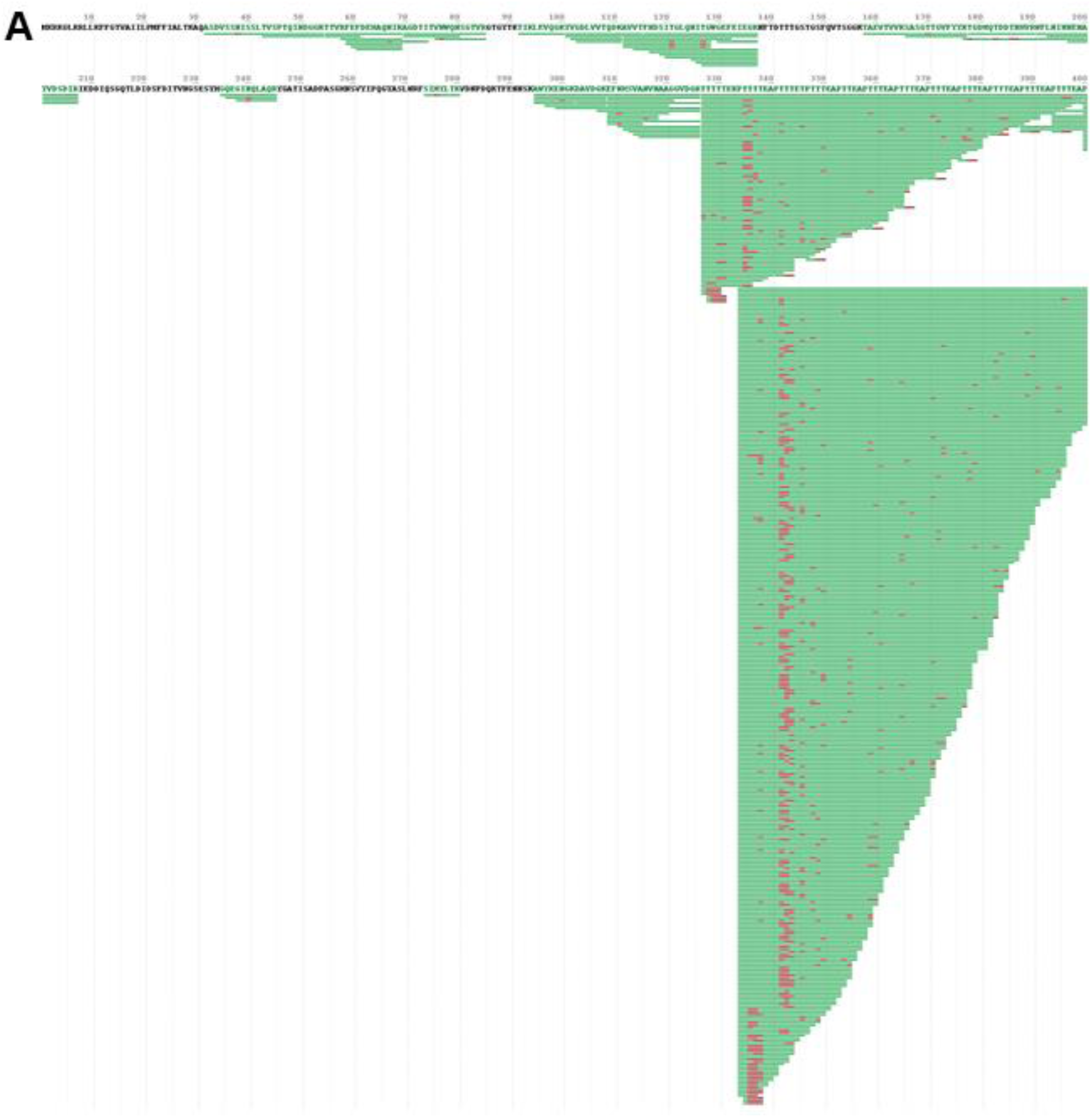
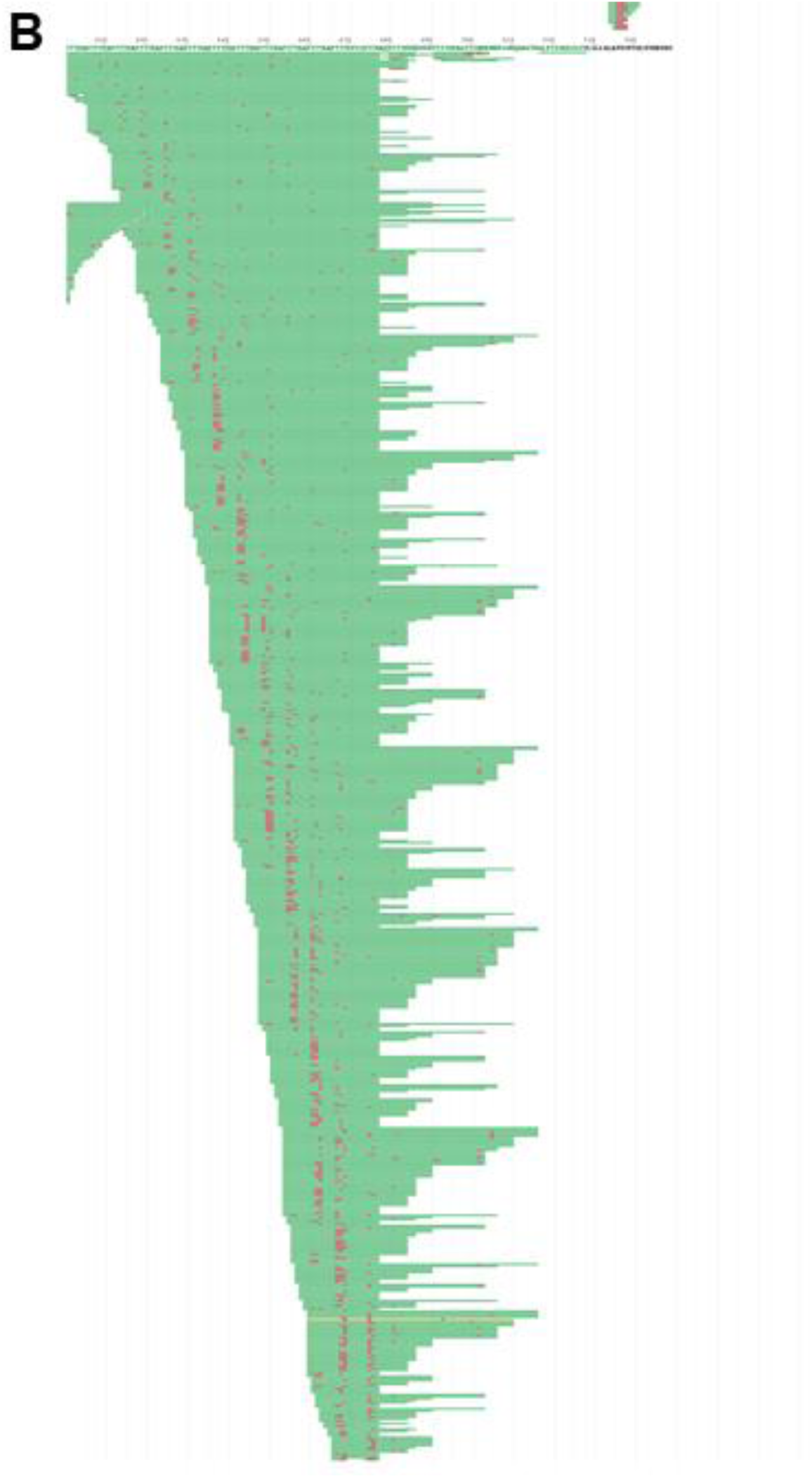
Phosphate coverage of Cnm from the *ΔpgfS* mutant. (A) N-terminal portion of full-length Cnm from the *ΔpgfS* mutant was subjected to phosphorylation mapping with Byonic. (B) C-terminal portion of Cnm, same map as panel A. Red markings at threonines are indications of phosphorylation. Other posttranslational modifications from processing, including oxidation on methionines and deamidation on asparagines, are also indicated. Partial cleavage (see methods) was used to obtain fragments in the threonine-rich repeat region yielding peptides with jagged ends. One of these phosphopeptides is illustrated in Figure 7.

**Figure S10.**
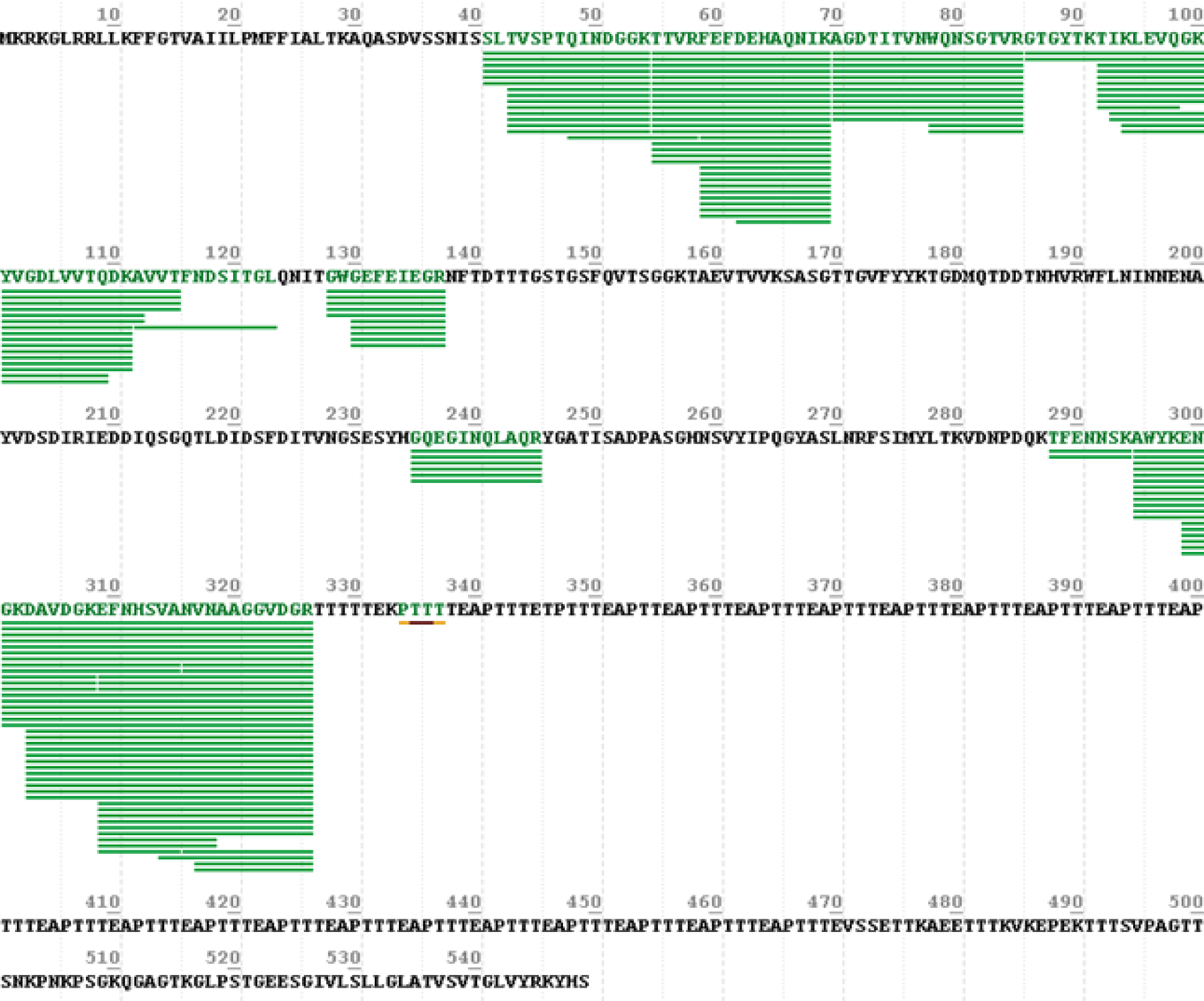
Phosphate coverage of Cnm from parental strain OMZ175. Mass spectrometry data were subjected to the same search with Byonic as Cnm derived from the *ΔpgfS* mutant. No indications of phosphorylation were found in this sample (as evidenced by the lack of red markings). The lack of coverage in the threonine-rich repeat region is due to the heavy glycosylation we observed in another study (Andresen *et al*., 2022a).

